# White matter plasticity and reading instruction: Widespread anatomical changes track the learning process

**DOI:** 10.1101/268979

**Authors:** Elizabeth Huber, Patrick M. Donnelly, Ariel Rokem, Jason D. Yeatman

## Abstract

White matter tissue properties correlate with children’s performance across domains ranging from reading, to math, to executive function. We use a longitudinal intervention design to examine experience-dependent growth in reading skills and white matter in a group of grade school aged, struggling readers. Diffusion MRI data were collected at regular intervals during an 8-week, intensive reading intervention. These measurements reveal large-scale changes throughout a collection of white matter tracts, in concert with growth in reading skill. Additionally, we identify tracts whose properties predict reading skill but remain fixed throughout the intervention, suggesting that some anatomical properties may stably predict the ease with which a child learns to read, while others dynamically reflect the effects of experience. These results underscore the importance of considering recent experience when interpreting cross-sectional anatomy-behavior correlations. Widespread changes throughout the white matter may be a hallmark of rapid plasticity associated with an intensive learning experience.

---

Skilled reading requires orchestration of a large cortical network, and individual differences in reading performance have been linked to the properties of white matter tracts connecting portions of this network specialized for processing visual, acoustic, and semantic features^1–9^. Although individual differences in white matter are thought to reflect the joint influence of genetics and experience^10–12^, white matter properties are often held to underlie variation in performance and to causally influence individual learning trajectories^13–16^. A number of recent studies, working within this framework, have identified features of the white matter that predict reading outcomes in dyslexia^17^, and reading-related skills, like phonological awareness, in prereading children^14,18,19^. The implication of these observations is that underlying anatomical differences may predestine certain individuals to struggle with learning to read. In this view, differences in white matter properties could be considered a reflection of intrinsic deficits, which might be relatively resistant to remediation, but which could plausibly be used for early identification of individuals in need of extra educational support^20^.

Successfully relating anatomical differences with behavioral outcomes requires an understanding of the timescale over which white matter tissue properties exhibit experience-dependent change, and the anatomical specificity of these effects. White matter plasticity, including activity-dependent myelination and oligodendrocyte proliferation, has been observed in animal models over the time-scale of days to weeks^21‒24^, and these effects coincide with changes in tissue properties measured non-invasively using diffusion-weighted magnetic resonance imaging (dMRI) in animals^25,26^. It has further been suggested that myelination may play a causal role in skill learning, since blocking the production of new myelinating oligodendrocytes inhibits motor skill development in mice^27^, implying that changes in white matter are critical to the learning process, rather than epiphenomenal. It is not clear whether similar effects occur in the context of human learning, particularly for a complex skill like reading, which is typically acquired with many hours of practice over a large developmental window. However, the studies cited above strongly suggest that learning should be accompanied by rapid, measurable changes in white matter. Further, a number of recent studies highlight the surprising malleability of human white matter in response to short-term training^28–31^, including training of reading and related skills^32–34^. This opens the possibility that correlations between white matter properties and behavior arise as temporary states within a highly plastic system that flexibly adapts to environmental demands. In this case, observed relationships between anatomy and behavior might be less stable than often presumed, given an appropriate change to the educational environment.

Here, we test whether controlled changes to a child’s educational environment induce changes in white matter tissue properties over the time-scale of weeks. Using a longitudinal intervention design, we track improvements in reading skills, and accompanying changes in white matter, in a group of grade-school aged, struggling readers during 8 weeks of intensive (4 hours each day, 5 days a week), one-on-one training in reading skills. We first examine learning effects within three tracts thought to carry signals critical for skilled reading^1–9,14,15,18,35–40^: the left arcuate fasciculus (AF), left inferior longitudinal fasciculus (ILF), and posterior callosal connections (CC). These pathways connect canonical reading-related regions within the ventral occipitotemporal (including the visual word form area, or VWFA^41–47^), superior temporal^48,49^, and inferior frontal cortex^50^, and hence, these tracts are considered to be part of the core circuitry for reading^37,46,47^. We find that the AF and ILF exhibit experience-dependent change within weeks the intervention onset, while tissue properties within the posterior CC remain fixed. Moreover, we illustrate the ambiguity of brain-behavior correlations measured in a dynamic system: As training rapidly alters an individual’s white matter and behavior, cross-sectional correlations between white matter properties and reading skills change substantially between measurement sessions. Meanwhile, CC white matter properties, which do not change during training, remain correlated with reading skill throughout the intervention. We therefore suggest that some anatomical properties may be stable predictors of the ease with which a child learns to read, while others dynamically reflect the effects of experience. These effects likely arise from distinct mechanism that cannot be distinguished by cross-sectional studies. Finally, we test the hypothesis that experience-dependent plasticity is anatomically localized to specific tracts. Contrary to this anatomical-specificity hypothesis, we find that educational experience alters a widespread system of white matter tracts in concert with reading skills. This system includes, but is not limited to, the core reading circuitry.

## Results

### Tracts connecting the core reading circuitry correlate with pre-intervention reading skill

We began by replicating previously reported correlations between reading skill and properties of the white matter tracts connecting key components of the reading circuitry^1–9,14,15,18,35–40^. To summarize individual differences in reading, we report Reading Skill, a composite score that incorporates our full battery of reading tests from the Woodcock-Johnson^51^ and TOWRE^52^ standardized assessments (see Methods for details, and **Supplementary Figure 1A**). To characterize the cross-sectional relationship between white matter and reading, we calculated simple, bivariate correlations between Reading Skill and each diffusion metric at the preintervention baseline session. As shown in **Figure 1**, pre-intervention (Session 1) measurements replicate previously reported correlations between reading scores and diffusion properties in the left arcuate, left ILF, and the CC: Correlations between MD and Reading Skill are positive both in the intervention group, and in the full sample, containing intervention and control subjects (**Figure 1**). The Reading Skill composite is a weighted sum of the individual reading tests, and similar effects are observed when examining correlations with the Woodcock Johnson (insert stats) and TOWRE (insert tests). Mirroring these effects, correlations between FA and reading are negative (**Supplementary Figure 2**). While several previous studies report a negative relationship between FA and reading in these pathways^3,4,8,38^, others report a positive relationship between FA and reading^1,2,7,35,53,54^. Thus, while properties of these pathways have consistently been shown to correlate with reading skill, the direction of this relationship is not consistent across studies, or tracts (see^55^). These inconsistencies may depend on factors like age, education, or SES, or may reflect the inherent ambiguity of dMRI metrics like FA, which can be influenced by a number of underlying biological phenomena with distinct, and potentially opposing, relationships to reading^38^.

**Figure 1.**
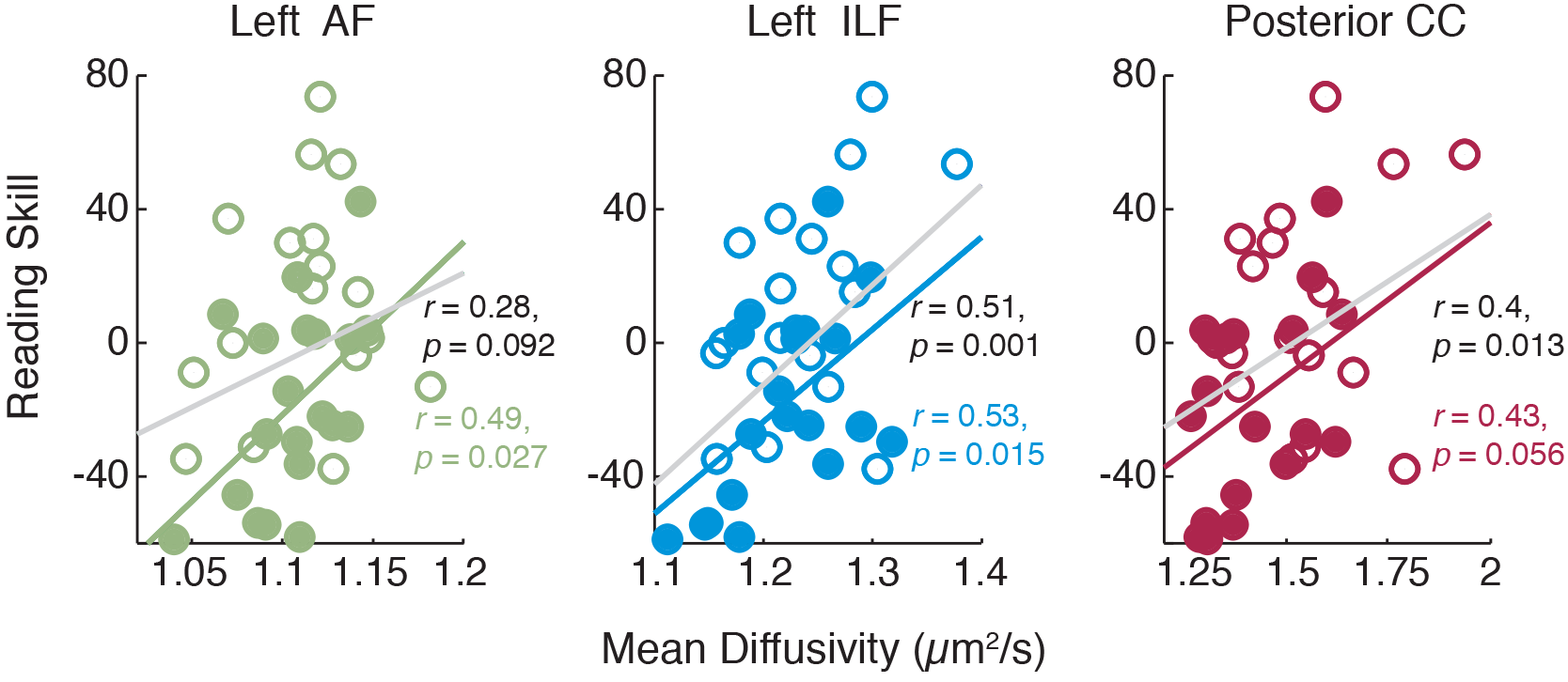
Pathways connecting the core reading circuitry correlate with pre-intervention Reading Skill. Tract average mean diffusivity (MD) is plotted as a function of pre-intervention (Session 1) reading skill. Best-fit lines plotted in gray give estimates for the full data set, while colored lines show fits for the intervention subjects, alone. Correlations for the intervention subjects are given in colored text below the value estimated for the full data set (in black).

In addition to the tracts chosen *a priori* for analysis, we examined several other tracts previously shown to correlate with reading scores, albeit less consistently across studies^9,56–58^, in a subsequent exploratory analysis. As shown in **Supplementary Table 1**, the left inferior frontal occipital fasciclus (IFOF) was also significantly correlated with reading skill (Bonferroni corrected *p* < 0.05), and a number of other tracts showed moderate, non-significant correlations. Finally, to test whether correlations were specific to Reading Skill, as opposed to general academic ability, we calculated correlations with math scores (Woodcock-Johnson Calculation and Math Facts Fluency) and found that none of the tracts that significantly correlated with reading (including the AF, ILF and callosal connections) correlated with math skills. Indeed, neither MD nor FA showed a significant relationship to math skills in any of the tracts chosen for analysis.

### Intensive intervention changes reading skills and white matter tissue properties

Reading skills improved substantially during the 8-week intervention period. Standard scores on the Woodcock-Johnson Basic Reading Composite, an untimed measure of reading accuracy, improved significantly over the course of the intervention (*F*(1,77) = 59.75, *p* < 10^−10^, linear mixed effect model with a fixed effect of intervention time, in hours, and a random effect of subject). After 8 weeks, the intervention-group mean was within one standard deviation of the population norm (100 +/− 15): pre vs. post intervention scores were 80.00 +/− 3.50 vs. 92.94 +/− 2.50). In line with these results, scores on the TOWRE Index, a timed measure of reading, improved substantially (*F*(1,77) = 53.69, *p* < 10^−9^), as did scores on the Woodcock-Johnson Reading Fluency subtest (*F*(1,76) = 36.042, *p* < 10^−7^). In contrast, we found no evidence for change in math skills during the intervention (Woodcock-Johnson Calculation Score, *F*(1,63) = 2.54, *p* = 0.12; Woodcock-Johnson Math Fact Fluency: *F*(68) = 1.87, *p* = 0.18), confirming that the intervention specifically affected reading skills. Additional details of these analyses are given in **Supplementary Figure 1**.

Growth in reading skill was specific to the intervention group, as indicated by a significant group (intervention versus control) by time (days) interaction for all reading, but not math, measures using a reading-skill-matched subgroup of the control subjects (n = 9). For this analysis, we substituted ‘days’ for ‘intervention hours’, to provide a meaningful index of time for both the intervention and control groups. For intervention subjects, ‘days’ were highly correlated with ‘intervention hours’, since testing sessions were scheduled at regularly spaced intervals (*r*(78) = 0.95, *p* < 001). For WJ Basic Reading Skills, we saw no significant effect of group (*F*(1,94) = 0.16, *p* = 0.68) or time (*F*(1,94) = 0.19, *p* = 0.67), but a significant group-by-time interaction (*F*(1,94) = 4.22, *p* = 0.042), indicating that growth in reading skills during the intervention period was specific to the intervention subjects. Similarly, for the TOWRE Index, we saw no significant effect of group (*F*(1,94) = 1.12, *p* = 0.29) or time (*F*(1,94) = 0.24, *p* = 0.63), but a significant group-by-time interaction (*F*(1,94) = 4.069, *p* = 0.047). For the WJ Calculation test, we saw a significant main effect of group (*F*(1,94) = 4.10, *p* = 0.046) but not time (*F*(1,94) = 0.31, *p* = 0.58), and no significant group-by-time interaction (*F*(1,94) = 1.13, *p* = 0.29), consistent with stability of this measure in both groups. Results for the full control sample (including both good and poor readers, n = 19) are given in **Supplementary Table 2**; this analysis shows that amongst the skilled reading control subjects, performance improved with repeated testing for the timed measures (TOWRE and Reading Fluency). In all control subjects, untimed measures (WJ Basic Reading) were stable, showing no change over 8 weeks. In other words, skilled readers benefitted slightly from repeated practice with the timed reading tests, while poor readers did not show any improvements with practice, and only showed an improvement in performance as a result of the intervention program.

To test whether changes in reading skill were accompanied by measurable changes in white matter structure, we first examined MD and FA as a function of intervention time (hours) within the set of white matter tracts considered to be crucial for skilled reading^1–9,14,15,18,35–39^, and which showed significant relationships with pre-intervention reading skill in the current sample: the left arcuate fasciculus (AF), left inferior longitudinal fasciculus (ILF), and posterior callosal connections (CC). Intervention driven tissue changes were evident within the AF and ILF, but not the CC: Specifically, mean diffusivity (MD) decreased as a function of intervention hours in the intervention within the left **AF** (*F*(1,77) = 8.46, *p* = 0.0047, linear mixed effect model with a fixed effect of intervention time, in hours, and a random effect of subject) and the left **ILF** (*F*(1,77) = 7.28, *p* = 0.0086), but not the **CC** (*F*(1,77) = 2.37, *p* = 0.13). Subject motion did not change over time (**Supplementary Figure 3**), and including subject motion as a covariate in the model did not change the results: MD decreased as a function of intervention hours within the left **AF** (*F*(1,76) = 10.48, *p* = 0.0018) and the left **ILF** (*F*(1,76) = 9.53, *p* = 0.0028), but not the **CC** (*F*(1,76) = 2.11, *p* = 0.15). The decline in mean diffusivity was accompanied by a linear increase in fractional anisotropy (FA) in the left **AF** (*F*(1,76) = 3.98, *p* = 0.050, fixed effect of intervention hours and a random effect of subject, with subject motion included as a covariate, as above) and the left **ILF** (*F*(1,76) = 8.82, *p* = 0.0040) but not the **CC**(*F*(1,76) = 0.24, *p* = 0.62). Finally, since changes in white matter properties could theoretically follow a nonlinear trajectory, we tested a model that included a quadratic term for each tract and parameter. For MD in each tract, the linear model outperformed the nonlinear model (evaluated using Bayesian Information Criteria, BIC^59,60^), and no significant nonlinear effects were observed: **AF** linear: *F*(1,76) = 8.72, *p* = 0.0041, **AF** quadratic: *F*(1,76) = 0.31, *p* = 0.58, **ILF** linear: *F*(1,76) = 7.53, *p* = 0.0076, **ILF** quadratic: *F*(1,76) = 0.33, *p* = 0.57, **CC** linear: *F*(1,76) = 3.083, *p* = 0.083, **CC** quadratic: *F*(1,76) = 3.90, *p* = 0.052. In contrast, we observed significant quadratic effects in FA for the left AF, only: **AF linear:** *F*(1,76) = 3.87, *p* = 0.053, **AF** quadratic: *F*(1,76) = 7.77, *p* = 0.0067, **ILF** linear: *F*(1,76) = 8.85, *p* = 0.0039, **ILF** quadratic: *F*(1,76) = 3.20, *p* = 0.078, **CC** linear: *F*(1,76) = 0.31, *p* = 0.58, **CC** quadratic: *F*(1,76) = 2.047, *p* = 0.16.

Like the reading outcomes reported above, intervention-driven changes in MD were specific to the intervention group, as indicated by a significant group (intervention versus control) by time (days) interaction. As above, we substitute ‘days’ for ‘intervention hours’, to give a meaningful predictor for both the intervention and control subjects. In the left **AF**, we found a significant main effect of group (*F*(1,125) = 7.047, *p* = 0.009) but not time (*F*(1,125) = 1.033, *p* = 0.31), and a significant group-by-time interaction (*F*(1,125) = 4.97, *p* = 0.028), consistent with a decrease in MD over time that was specific to the intervention subjects. Similarly, in the **ILF**, we saw a significant main effect of group (*F*(1,125) = 10.29, *p* = 0.0017) but not time (*F*(1,125) = 3.72, *p* = 0.056), and a significant group-by-time interaction (*F*(1,125) = 9.53, *p* = 0.0025). In the **CC**, we saw a significant main effect of group (*F*(1,125) = 6.69, *p* = 0.011) but not time (*F*(1,125) = 0.90 *p* = 0.34), and no significant group-by-time interaction (*F*(1,125) = 0.027, *p* = 0.87), consistent with the stability of MD values in this tract in all subjects. For FA, we observed a different pattern of results: In the **AF**, we saw no significant main effect of group (*F*(1,125) = 0.31, p = 0.58) or time (*F*(1,125) = 0.055, p = 0.82), and no significant group-by-time interaction (*F*(1,125) = 0.36, p = 0.55). In the **ILF**, we saw no significant main effect of group (*F*(1,125) = 0.0015, *p* = 0.97) or time (*F*(1,125) = 1.93, *p* = 0.17), and no significant group-by-time interaction (*F*(1,125) = 0.15, *p* = 0.70). In the **CC**, we saw no significant main effect of group (*F*(1,125) = 0.23, *p* = 0.63) or time (*F*(1,125) = 0.86, *p* = 0.36), and no significant group-by-time interaction (*F*(1,125) = 0.35, *p* = 0.56). As shown in **Supplementary Table 3**, the group-by-time interaction approached significance for the quadratic term for FA in the left AF and ILF, but not for MD in the AF or ILF, or for either parameter in the CC.

Given the observed non-linearity of intervention-driven effects in FA, we opted to use ‘session number’ as a categorical predictor in the analysis to follow, since this approach summarizes session-to-session differences from baseline, without imposing a shape on the trajectory of change. Sessions were systematically spaced over time, and this timing was consistent across subjects; hence ‘session’ was highly correlated with ‘days’ (*r*(127) = 0.97, p < 0.001). As shown in Figure 2, both the left AF and ILF showed clear intervention-driven changes in both MD and FA. Within the intervention group, significant changes in tissue properties emerged in the first post-baseline measurement session, after just 46.05 hours (SD = 14.88) of intervention, over the course of 2-3 weeks. In line with the results reported above for the continuous predictor (days), we observed a group-by-session interaction for MD in the **AF** (no main effect of session, *F*(1,67) = 2.12, *p* = 0.15, or group, *F*(1,67) = 0.58, *p* = 0.45, session-by-group interaction, *F*(1,67) = 7.75, *p* = 0.0070), and the **ILF** (no main effect of session, *F*(1,67) = 1.77, *p* = 0.19, or group, *F*(1,67) = 0.044, *p* = 0.83, session-by-group interaction, *F*(1,67) = 6.91, *p* = 0.011) but not the **CC** (no main effect of session, *F*(1,67) = 1.029, *p* = 0.31, main effect of group, *F*(1,67) = 5.99, *p* = 0.017, no session-by-group interaction, *F*(1,67) = 0.62, *p* = 0.44), and for FA in the **ILF** (no main effect of session, *F*(1,67) = 0.65, *p* = 0.42, or group, *F*(1,67) = 0.60, *p* = 0.44, session-by-group interaction, *F*(1,67) = 6.45, *p* = 0.013), but not the **AF** (no main effect of session, *F*(1,67) = 0.0057, *p* = 0.94, or group, *F*(1,67) = 1.57, *p* = 0.21, no session-by-group interaction, *F*(1,67) = 2.85, *p* = 0.096) or **CC** (no main effect of session, *F*(1,67) = 0.26, *p* = 0.61, or group, *F*(1,67) = 0.14, *p* = 0.71, no session-by-group interaction, *F*(1,67) = 2.38, *p* = 0.13). An exploratory analysis of this same session-by-group interaction for all available tracts is given in **Supplementary Figure 10**. Finally, to ensure that the interaction was not driven by differences in the stability of our measurements in good vs. poor readers, given that the control group included both typical readers and subjects with dyslexia, we repeated the above analysis with baseline Reading Skill included as a covariate in the model. We obtained the same results for the group-by-session interaction in all cases (**AF**: MD, *F*(1,65) = 7.72, *p* = 0.0071; FA, *F*(1,65) = 2.86, *p* = 0.095; **ILF**: MD, *F*(1,65) = 8.37, *p* = 0.0052; FA, *F*(1,65) = 6.71, *p* = 0.012; **CC**: MD, *F*(1,65) = 0.63, *p* = 0.43; FA, *F*(1,65) = 2.42, *p* = 0.12).

**Figure 2.**
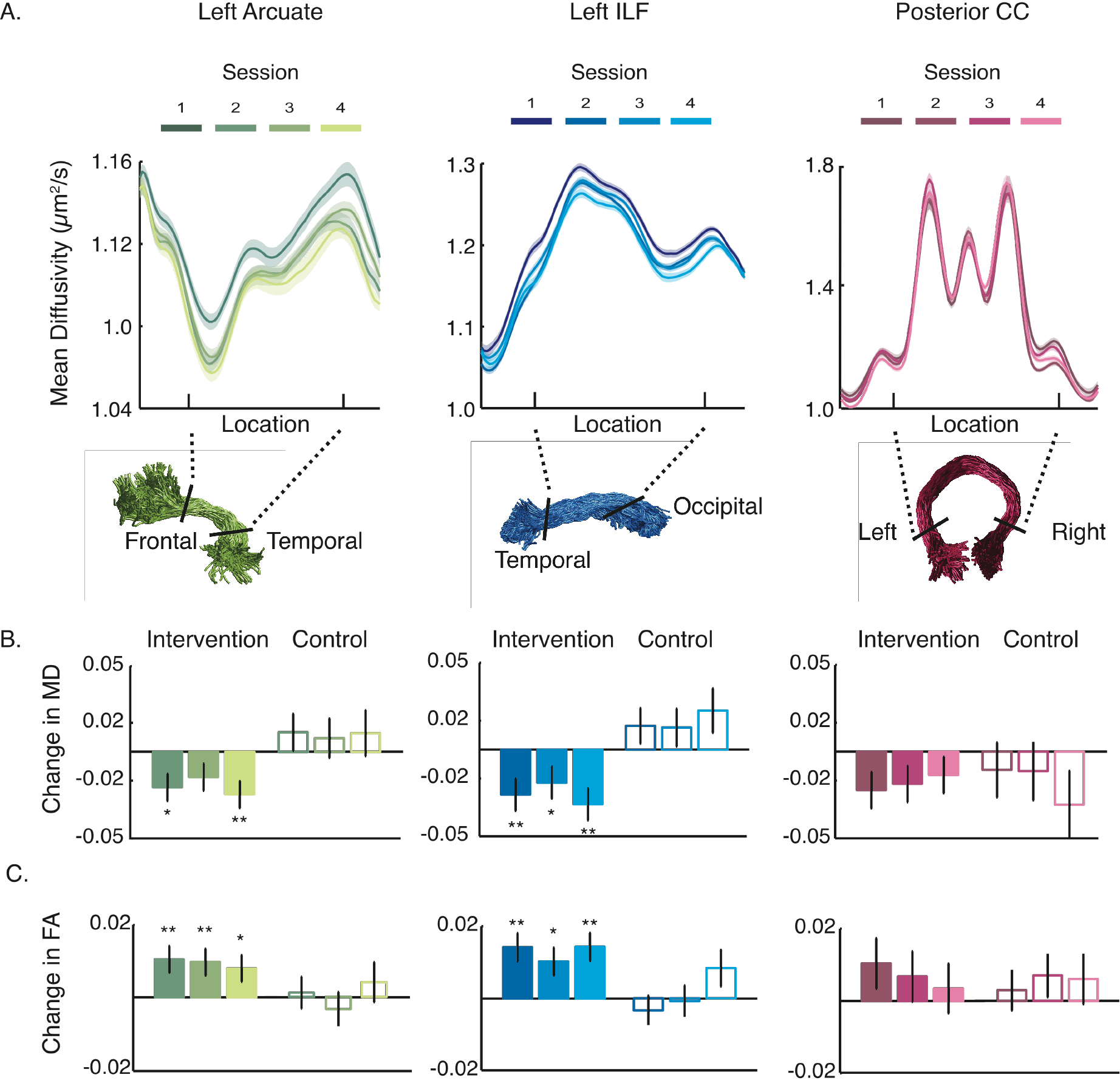
Change versus stability in Tract Profiles during reading intervention. (A) Mean diffusivity values were mapped onto each of 100 evenly spaced nodes spanning termination points at the gray-white matter boundary to create a ‘Tract Profile’ (see Methods and^61^ for additional details of this analysis). For visualization purposes, the middle 80 nodes are plotted. Each line represents the group average mean diffusivity (MD) across subjects, measured at four time-points: pre-intervention (Session 1), after ~2.5 weeks of intervention (Session 2), after ~5 weeks of intervention (Session 3), and after 8 weeks of intervention (Session 4). Shaded error bars give ±1 standard error of the mean. Color values indicate session, ranging from darkest (Session 1) to brightest (Session 4) for each tract. The x-axis shows the location where each tract was clipped prior to analysis (corresponding to black boundary lines in renderings, above). Tract renderings are shown for an example subject. The middle 60% (bounded by black lines) of each tract was analyzed in (B-C), to avoid partial volume effects that occur at endpoints of the tract, where it enters cortex. Both the AF and ILF, but not the posterior callosal connections, show a systematic decrease in MD over the course of intervention. (B-C) Bars show model predicted change (coefficients and standard errors from mixed effects model) in MD (B) and FA (C) for each session. Bar heights represent the magnitude of change observed in that session, relative to Session 1 (pre-intervention) baseline, as determined by the mixed effects model. As described in the main text, both the arcuate fasciculus (AF) and inferior longitudinal fasciculus (ILF) showed significant change between sessions for the intervention group (filled bars), but not the control group (unfilled bars). Asterisks indicate a significant decrease in MD (B) or increase in FA (C) for each session relative to the pre-intervention baseline at a Bonferroni corrected *p* < 0.05 (*) and *p* < 0.01 (**).

### Reading intervention does not ‘normalize’ differences in the white matter

One possible interpretation of group differences in MD and FA between good and poor readers is that these differences reflect abnormal tissue properties in poor readers. In that case, one could expect that remediation of reading difficulties might be associated with a “normalization” of deficits in white matter structure. However, we find that intervention driven changes in white matter do not follow the trajectory predicted by a normalization account. **Figure 3** shows changes in MD and reading scores for the intervention group, relative to the subset of non-intervention controls who had reading skills in the typical range. We defined ‘Typical Readers’ as Control Group subjects with timed (TOWRE Index) and untimed (WJ Basic Reading Score) reading accuracy within a standard deviation of the population mean (at or above 85 on both measures). For the intervention group, we plot changes in both WJ Basic Reading and the TOWRE Index (rather than composite Reading Skill), in order to situate the cross-sectional and intervention-driven effects relative to an age-normed, population mean. After completing the intervention, tissue properties in the intervention subjects were not more similar to the typical reading controls, despite a substantial improvement in reading skills. As diffusion properties such as MD are influenced by multiple biological sources, this finding indicates that short-term plasticity is likely to reflect a different biological mechanism than the group differences reported here and in other studies. Further, the short-term, experience-dependent changes in the white matter were larger (Cohen’s *d* = 0.75 for the **AF**, and *d* = 0.66 for the **ILF**) than typical group difference reported in the literature^8,19,53^, and the group differences observed here (*d* = 0.53 for the **AF** and *d* = 0.59 for the **ILF**). These results demonstrate that the effects of recent experience on measured tissue properties may equal or exceed effects due to intrinsic or long-term maturational factors, suggesting that group differences measured in cross-sectional studies may, in some cases, be driven by systematic differences in environmental influences between groups.

**Figure 3.**
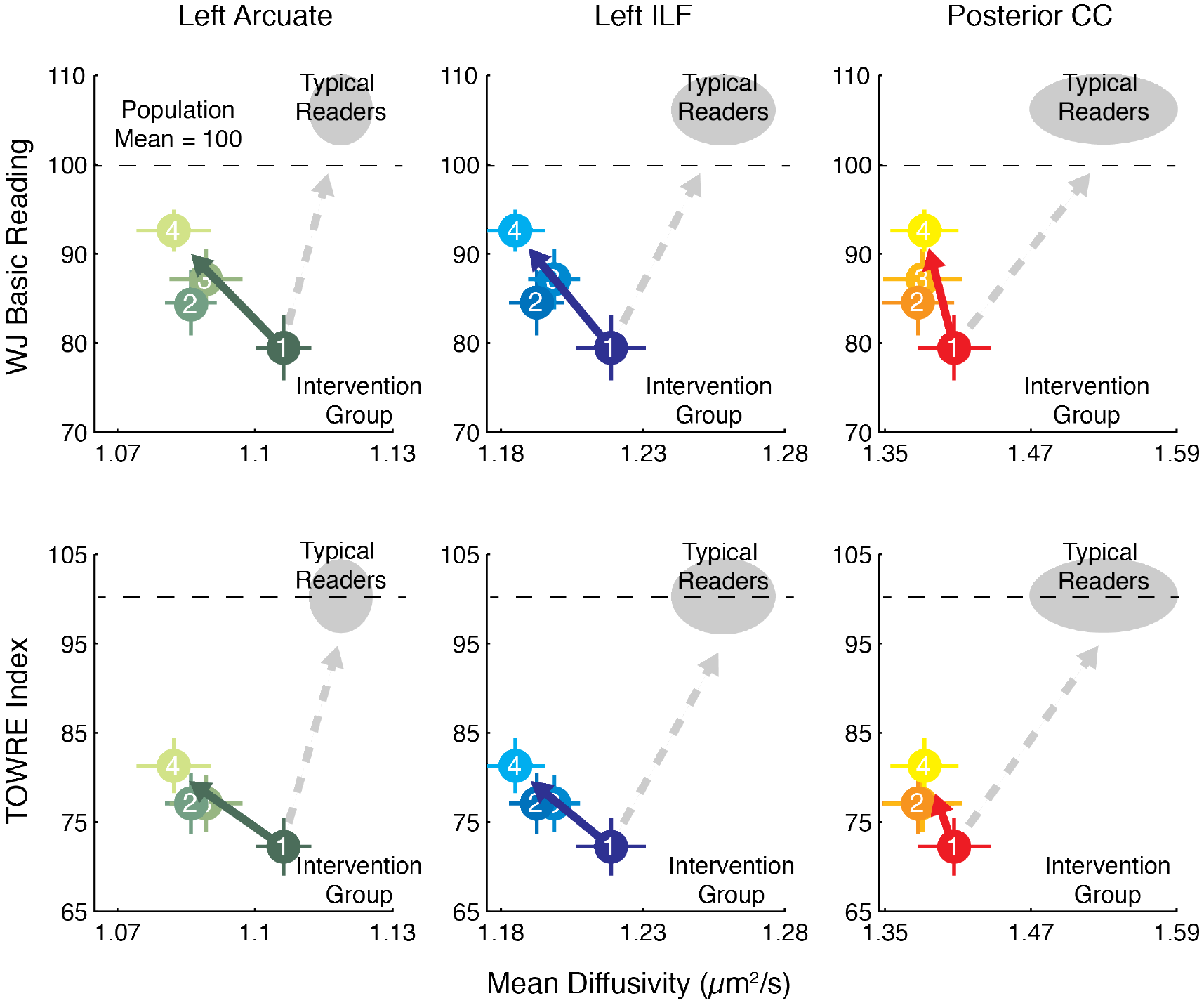
Reading intervention does not ‘normalize’ differences in the white matter. Reading skills are plotted as function of mean diffusivity for each session (1-4) for the left arcuate, ILF, and posterior callosal tracts in the intervention group. The gray ellipse in each panel shows the mean and standard error for the subset non-intervention control subjects with reading skills in the typical range (poor reading controls were excluded, leaving n =10 typical reading controls). The dashed gray arrow shows the expected trajectory for MD values if the intervention group were to become more similar to the typical reading controls in terms of both reading skills and MD. In contrast, the true trajectory of change in plotted as a colored arrow in each panel. The intervention group includes some readers with only moderate reading impairments (and, therefore, higher MD values), and so the group difference in pre-intervention scores is less than would be expected for a group of good vs. poor readers.

### Anatomy-behavior correlations depend on recent experience for the arcuate and ILF

Over the course of the intervention, only the posterior callosal connections retained their relationship to Reading Skill. In contrast, as MD values declined in the AF and ILF, the instantaneous, cross-sectional correlation between reading and MD changed between sessions, as indicated by a significant interaction between MD and session in predicting Reading Skill for the intervention group (linear mixed effects model predicting Reading Skill from MD, session, and their interaction, with a random effect of subject, see **Figure 4**). For MD in both the AF and ILF, but not the CC, this interaction was significant (main effect of MD in **AF**, *F*(1,71) = 4.59, *p* = 0.036, main effect of session *F*(3,71) = 28.048, *p* < 10“^−11^, session-by-MD interaction, *F*(3,71) = 2.95, p = 0.039; main effect of MD in **ILF**, *F*(1,71) = 3.97, p = 0.050, main effect of session *F*(3,71) = 28.53*,p* < 10^−11^, session-by-MD interaction, *F*(3,71) = 3.56, p = 0.018; main effect of MD in **CC**, *F*(1,71) = 1.56, *p* = 0.22, main effect of session *F*(3,71) = 27.19, *p* < 10^−11^, session-by-MD interaction, *F*(3,71) = 0.64, p = 0.59). In the AF, this effect was also significant for FA (main effect of FA in **AF,** *F*(1,71) = 8.48, p = 0.0047, main effect of session *F*(3,71) = 31.91, *p* < 10^−13^, session-by-FA interaction, *F*(3,71) = 4.28, p = 0.0078; main effect of FA in **ILF**, *F*(1,71) = 7.44, p = 0.0080, main effect of session *F*(3,71) = 30.99, *p* < 10^−13^, session-by-FA interaction, *F*(3,73) = 1.81, *p* = 0.15; main effect of FA in **CC**, *F*(1,71) = 9.43,*p* = 0.003, main effect of session *F*(3,71) = 32.60,*p* < 10^−12^, session-by-MD interaction, *F*(3,71) = 2.077, *p* = 0.11). Importantly, changes in both the strength and the sign (positive versus negative) of observed correlations could not be attributed simply to session-by-session changes in variance of reading skills or white matter. As shown in **Supplementary Table 4,** there was no statistical difference in variance across sessions (indeed, variances were nearly matched; see also **Figure 3,** which plots means and errors for each session). Therefore, changing anatomy-behavior correlations were not driven by differences in relative variance over time, and instead reflect learning related dynamics in the reading and white matter measures.

Finally, we found no evidence for changing anatomy-behavior correlations in the group of children who were not enrolled in the intervention (**AF:** MD: *F*(3,41) = 0.75, *p* = 0.53, FA: *F*(3,41) = 0.12, *p* = 0.95; **ILF**: MD: *F*(3,41) = 1.36, *p* = 0.27, FA: *F*(3,41) = 1.70, *p* = 0.18; **CC**: MD: *F*(3,41) = 0.55, *p* = 0.65, FA: *F*(3,41) = 0.97, *p* = 0.42). This is consistent with the stability of diffusion properties in this group, and supports the notion that the significant interaction for the intervention subjects did not arise due to differences in measurement noise over time. Finally, to rule out the possibility that systematic differences in head motion might influence anatomy-behavior correlations (e.g., children with lower reading scores might move more in the scanner than children with higher reading scores), we calculated the correlation between head motion and Reading Skill. Motion and Reading Skill were unrelated (*r*(97) = 0.13, *p* = 0.19).

**Figure 4.**
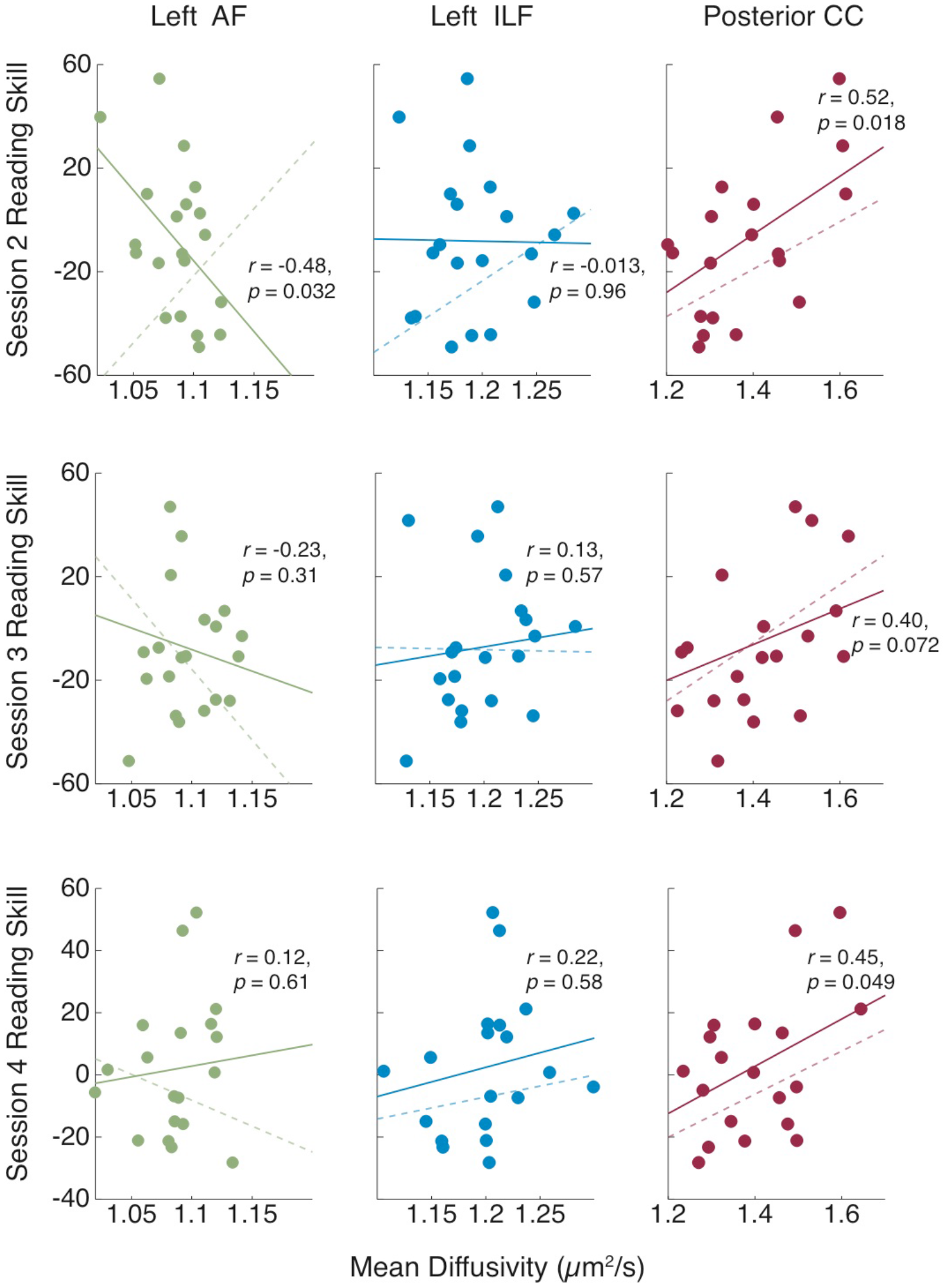
Correlations between white matter properties and reading skill change during learning. Plots show the cross-sectional correlation for Sessions 2-4 (top to bottom) for each tract for the intervention group. A solid best-fit line is plotted for each session. A dashed line in each panel represents the best-fit line for the preceding session (Session n - 1) to illustrate the session-to-session changes. For the AF and ILF, correlations change in size and/or direction, demonstrating that anatomy-behavior relationships can depend on recent educational experience.

### White matter properties of the AF and ILF change in concert and track individual learning

The AF and ILF connect distinct components of the reading circuitry and are thought to carry signals that contribute uniquely to the reading process ^37,47,55,62^. Therefore, a reading intervention might affect these tracts differently, prompting changes that reflect independent biological processes unfolding with different time-courses, and reflecting different aspects of learning. To address this possibility, we asked whether changes in the AF and ILF occur in synchrony in the intervention group. If wholly independent mechanisms were driving growth in both tracts, we would not expect to see similar time-courses of growth for the AF and ILF within subjects. Alternatively, if changes within the AF and ILF reflect a common biological mechanism operating over a large anatomical scale, then time-courses of growth should be correlated within subjects.

To address these questions, we fit a linear mixed effects model to all intervention subjects’ mean-centered diffusion measurements over all time points. This allowed us to quantify the similarity between AF and ILF longitudinal growth trajectories while excluding between-subject differences in baseline diffusion properties^63^. Results for complementary analysis, examining diffusion measurements relative to a pre-intervention baseline, is given in **Supplementary Table 5.** As shown in **Figure 5**, the time-courses of change in the AF and ILF were highly correlated for both MD and FA (MD: *r* = 0.86,*p* < 0.001; FA: *r* = 0.50, *p* = 0.021), implying that, within each individual, white matter growth trajectories were tightly coupled for these two tracts. We then fit the same model for time-lagged versions of each tract’s time-course, to test whether these regions changed in synchrony. If growth in one tract were to precede growth in the other, this would imply a distinct and more gradual process occurring in the second tract, or a possible causal relationship. In that case, the time-courses should be better predicted by time-lagged versions each other. However, we failed to detect a significant correlation at any non-zero lag, suggesting that these tracts change in concert as a function of experience in the reading intervention program.

Finally, to examine the relative timing of white matter changes in relation to learning, we performed the same cross-correlation analysis with the reading scores: Each intervention subject’s reading scores were mean-centered to remove inter-subject differences in baseline reading ability, and a linear mixed effects model was fit to shifted (lag = −1 and lag = 1) and un-shifted (lag = 0) versions of the time-courses. Time courses of MD, but not FA, were significantly correlated with time-courses of Reading Skill only at lag = 0 (MD: *r* = −0.30 Arcuate, *p* = 0.0069; *r* = −0.30 ILF, *p* = 0.0061), demonstrating that, within a subject, the time-course of white matter plasticity tracked the time-course of learning. For MD, we again found that the growth trajectories were best fit by the un-shifted time courses, suggesting that white matter changes are coupled to reading experience and, therefore, track improvements in Reading Skill. In the control group, no tracts showed a significant relationship to reading skill at any lag (shown for lag = 0 in **Supplementary Table 6**), consistent with the stability of both reading and white matter properties in control subjects.

**Figure 5.**
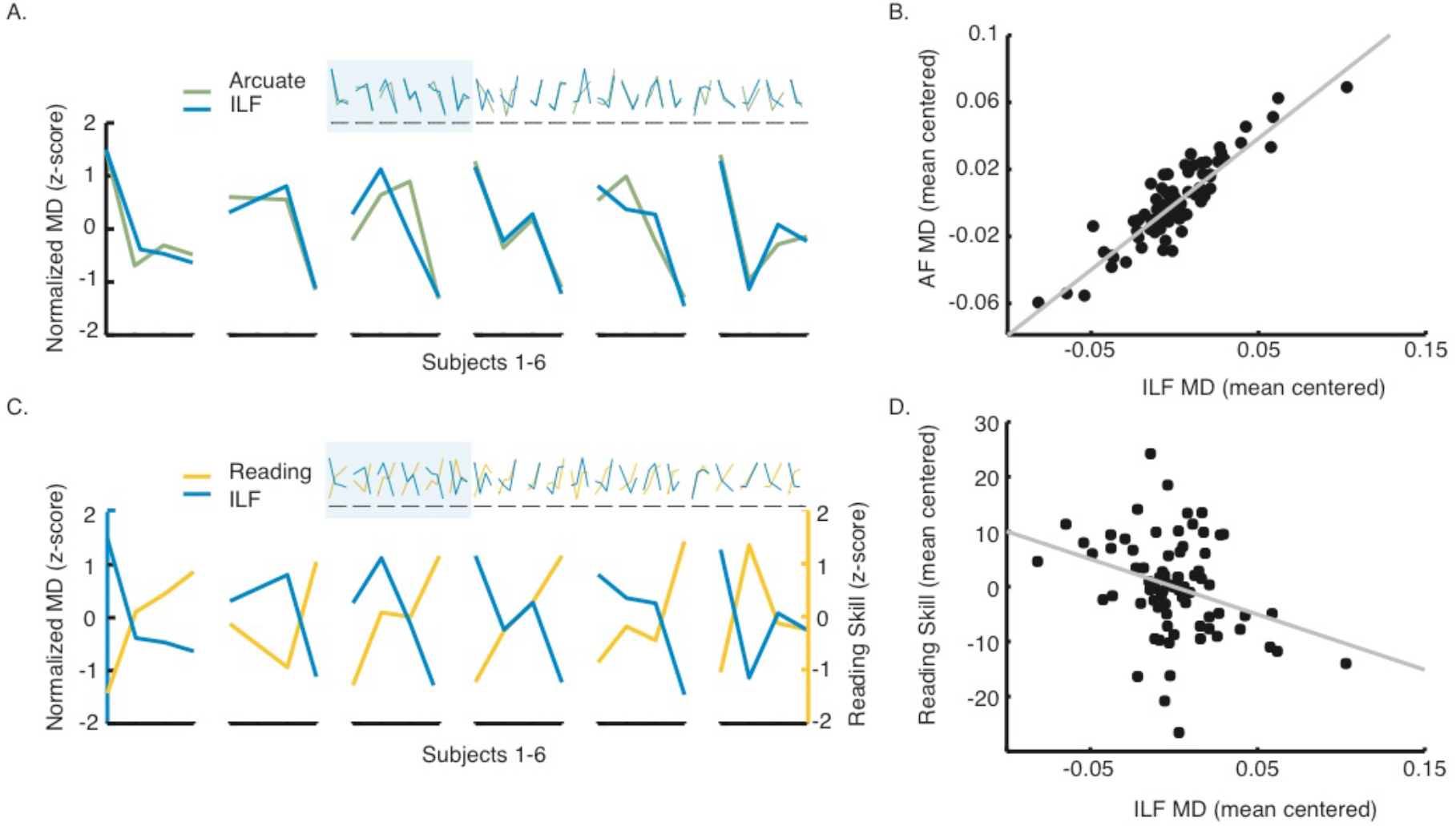
Analysis of individual growth trajectories in the AF and ILF. (A) Left Arcuate and ILF mean diffusivity (MD) time courses are correlated across individual subjects. For visualization, standardized MD values are plotted for each tract at each available time point for all subjects. Individual time-courses are enlarged for a set of 6 example subjects to show greater detail. Individual AF and ILF time-courses were positively correlated, and a cross-correlation analysis failed to detect a significant correlation at any nonzero lag, consistent with the interpretation that growth occurs in concert across tracts. (B) MD values were mean-centered for each subject, thus representing each subject’s time-course of MD changes as modulations around their mean. Mean-centered values at each available time point are plotted for the AF vs. ILF showing tight correspondence between changes in these two tracts. (C) The time-course of change in Reading Skill varies across subjects, and is negatively related to individual white matter time-courses: Within a subject, as MD decreases in the ILF, Reading Skill increase. For visualization, standardized Reading Skill and MD values are plotted for the ILF at each available time point for all subjects. As in panel (A), time-courses are enlarged for a set of 6 example subjects to show individual subject trajectories in greater detail. (D) Mean-centered MD values at each available time point for the ILF correlate with mean-centered Reading Skill assessed at each time point, demonstrating the relationship between time-courses of MD and Reading Skill change. The scatter plots in panel B and D also make it clear that the time-course of plasticity is more tightly coupled across tracts than it is to behavior. Hence, even though there is a statistically significant relationship between the time-course of white matter and behavioral changes, there is also un-explained variance that is likely to be related to aspects of the intervention environment that do not directly impact behavior.

Since a substantial proportion of the total changes in MD occurred during the first 2 weeks of intervention, we also examined the relationship between reading and white matter changes during this interval by correlating Session 2 vs. Session 1 difference scores. Individual differences in the magnitude of session 2-1 MD change were not significantly correlated with the magnitude of reading score change. As shown in **Supplementary Table 7**, we observed a trend for both raw and standardized reading scores. Since this analysis only includes half of the data, we cannot ascertain whether the result represents the absence of a relationship at this short timescale, or the lack of statistical power.

### Reading intervention prompts distributed changes in white matter

White-matter growth rates were highly correlated for two tracts considered to be critical for skilled reading, meaning that a subject showing rapid, intervention-driven growth in the AF also shows considerable growth in the ILF. However, changes in mean diffusivity and fractional anisotropy were not limited to the connections of the core reading circuitry; instead, we observed significant change throughout a collection of tracts, extending beyond our *a priori* hypothesis. **Figure 6a** models growth in MD as a linear function of the number of intervention hours, and we use a conservative Bonferroni correction in this exploratory analysis. In the intervention group, 12 out of 18 tracts showed significant (Bonferroni corrected) change. None of the 18 tracts showed significant change in either MD or FA in the control group. Further, as shown in **Table 1**, multiple tracts showed a significant relationship to changes in reading skill, including, but not limited to, the core circuitry for reading. (See **Supplementary Table 8** for a complementary analysis relating FA and Reading Skill.) Therefore, learning effects are not specific to tracts that are considered to be the core circuitry for reading, and intervention-driven changes are evident in an extensive collection of white matter tracts.

**Table 1.** White matter properties track changes in Reading Skill. Cells show p-values based on a mixed linear model predicting session-to-session changes Reading Skill composite score, Woodcock-Johnson Basic Reading scores, and TOWRE index scores, and Reading Fluency from changes in mean diffusivity (MD) at each time point during the intervention. Pearson correlations between mean-centered MD and mean-centered reading score are provided as an index of effect size. Tracts that predict changes in readings scores at a Bonferroni corrected *p* < 0.05 are highlighted in bold italic.

| Tract (MD) | Reading Skill Composite | Basic Reading Score | TOWRE Index | Reading Fluency |
| --- | --- | --- | --- | --- |
| Left Thalamic Radiation | $r=-0.025, p=0.026$ | $r=-0.16, p=0.17$ | **$r=-0.38, p=0.00056$** | $r=-0.15, p=0.18$ |
| Right Thalamic Radiation | $r=-0.31, p=0.0047$ | $r=-0.26, p=0.02$ | **$r=-0.37, p=0.00067$** | $r=-0.15, p=0.18$ |
| Left Corticospinal | **$r=-0.39, p=0.0003$** | $r=-0.32, p=0.0040$ | **$r=-0.39, p=0.00028$** | $r=-0.26, p=0.022$ |
| Right Corticospinal | **$r=-0.34, p=0.0019$** | $r=-0.27, p=0.014$ | **$r=-0.40, p=0.00025$** | $r=-0.21, p=0.062$ |
| Left Cingulum Cingulate | $r=-0.19, p=0.083$ | $r=-0.088, p=0.44$ | **$r=-0.34, p=0.0023$** | $r=-0.11, p=0.31$ |
| Right Cingulum Cingulate | $r=-0.12, p=0.29$ | $r=-0.13, p=0.23$ | $r=-0.11, p=0.31$ | $r=-0.050, p=0.66$ |
| Callosum Forceps Major | $r=-0.23, p=0.040$ | $r=-0.19, p=0.087$ | $r=-0.21, p=0.066$ | $r=-0.18, p=0.10$ |
| Callosum Forceps Minor | $r=0.041, p=0.72$ | $r=0.086, p=0.44$ | $r=-0.092, p=0.41$ | $r=0.077, p=0.50$ |
| Left IFOF | **$r=-0.33, p=0.0024$** | $r=-0.24, p=0.035$ | **$r=-0.42, p=0.00011$** | $r=-0.14, p=0.22$ |
| Right IFOF | $r=-0.28, p=0.013$ | $r=-0.25, p=0.024$ | $r=-0.25, p=0.024$ | $r=-0.089, p=0.43$ |
| Left ILF | $r=-0.30, p=0.0061$ | $r=-0.19, p=0.087$ | **$r=-0.40, p=0.00021$** | $r=-0.10, p=0.37$ |
| Right ILF | $r=-0.26, p=0.019$ | $r=-0.21, p=0.058$ | $r=-0.28, p=0.012$ | $r=-0.054, p=0.63$ |
| Left SLF | $r=-0.25, p=0.026$ | $r=-0.19, p=0.093$ | $r=-0.32, p=0.0037$ | $r=-0.089, p=0.43$ |
| Right SLF | $r=-0.25, p=0.022$ | $r=-0.20, p=0.077$ | $r=-0.31, p=0.0047$ | $r=-0.13, p=0.26$ |
| Left Uncinate | $r=-0.29, p=0.0081$ | $r=-0.24, p=0.029$ | $r=-0.31, p=0.0044$ | $r=-0.16, p=0.17$ |
| Right Uncinate | $r=0.037, p=0.74$ | $r=0.066, p=0.56$ | $r=0.0051, p=0.96$ | $r=0.17, p=0.13$ |
| Left Arcuate | $r=-0.30, p=0.0069$ | $r=-0.21, p=0.064$ | **$r=-0.40, p=0.00022$** | $r=-0.14, p=0.22$ |
| Right Arcuate | $r=-0.29, p=0.0082$ | $r=-0.24, p=0.030$ | **$r=-0.35, p=0.0015$** | $r=-0.15, p=0.18$ |

Given that intervention effects appear to be spatially widespread, and that changes within two key tracts, the AF and ILF, are tightly coupled, we next examined the correlation structure across the full collection of tracts showing intervention-driven growth. Specifically, we tested whether growth rates are solely coupled within the AF and ILF, versus a larger collection of tracts. To that end, we fit linear growth rates (change in MD or FA as a function of hours of intervention) to each subject’s data for the 18 tracts and then computed the correlation between growth rates across each pair of tracts. To assess the suitability of a linear model, we used Bayesian Information Criteria (BIC)^59,60^ to evaluate the linear model relative to two non-linear models, one with a quadratic and one with an additional cubic component. In all tracts with significant intervention-driven effects, the linear model outperformed both the quadratic and cubic models.

**Figure 6b** shows the correlation between linear growth rates of pairs of tracts across individuals. The ordering of the tracts was determined according to a hierarchical clustering of these correlation coefficients. This analysis revealed that many tracts show highly correlated intervention-driven changes (r > 0.7) and identified a cluster containing many of the cortical association tracts (the left and right ILF, SLF, IFOF, and arcuate, as well as the left uncinate and left corticospinal tracts) which all changed in concert. In addition, we identified a separate cluster of tracts whose properties change during the intervention, but with independent growth rates. For example, highly significant growth rates are observed bilaterally in the thalamic radiation, but these growth rates are not correlated with growth measured in the left arcuate (**Figure 6c**). Accordingly, these tracts are assigned to distinct clusters. We suggest that changes within these distinct clusters may reflect distinct biological mechanisms. A complementary analysis of FA is provided in **Supplementary Figure 2**, and identifies a consistent clustering of the tracts.

**Figure 6.**
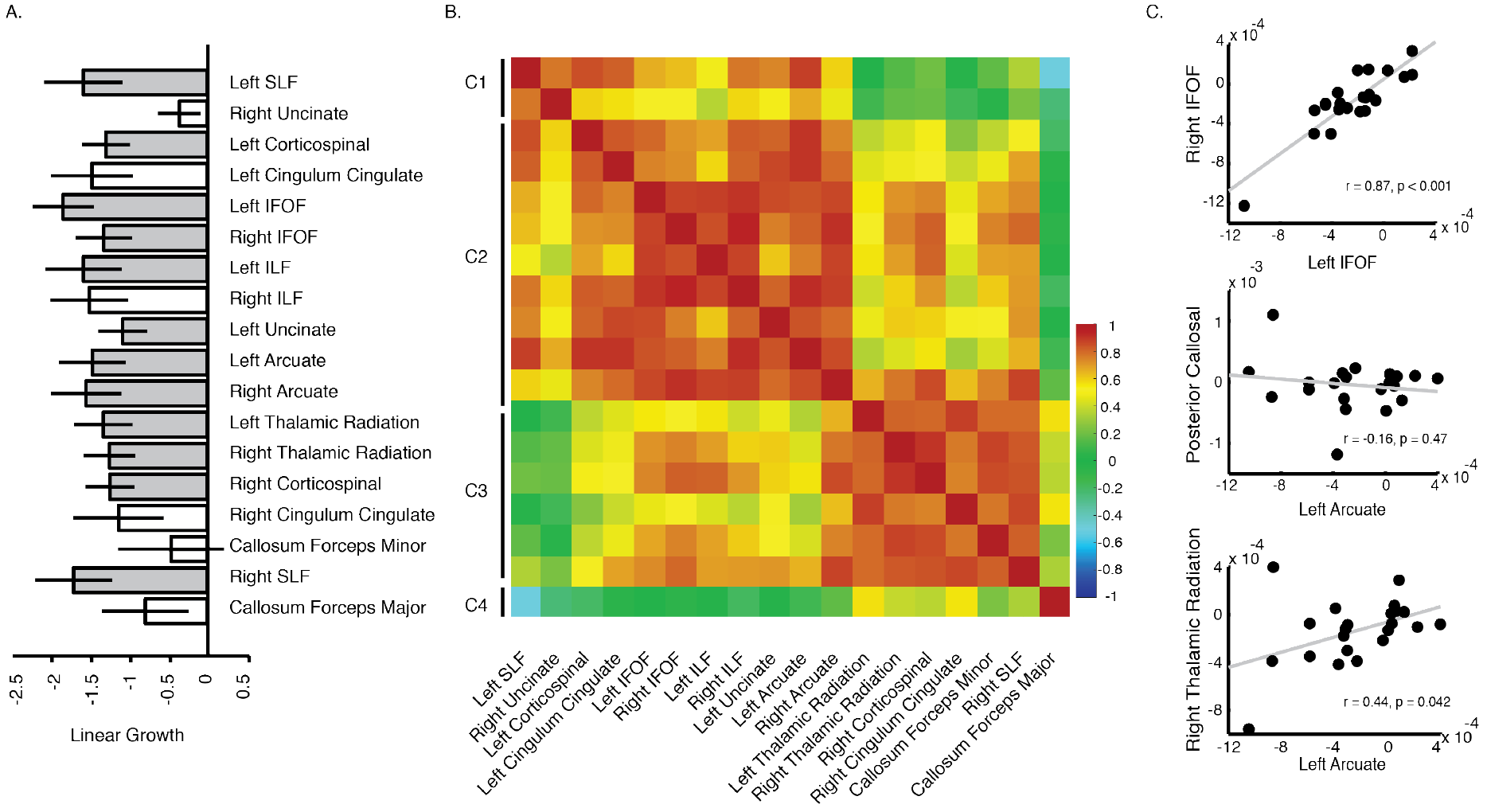
Reading intervention causes distributed changes in the white matter. (A) Change in MD as a function of intervention time (in hours) for 18 tracts. Tracts showing significant change (Bonferroni corrected *p* < 0.05) are indicated as gray filled bars. (B) Hierarchical clustering based on the correlations between linear growth rates. The heat map represents Pearson correlations between linear growth rates for pairs of tracts across individuals. The matrix is sorted according to hierarchical clustering of these correlation coefficients. (C) Scatter plots of individual growth rates for three pairs of tracts: left vs. right IFOF, AF vs. CC, and AF vs. right thalamic radiation.

## Discussion

Intensive reading training causes rapid changes in tissue properties within the left arcuate and inferior longitudinal fasciculus, two tracts considered crucial for skilled reading. However, the effects of intervention are not limited to these regions. Instead, we find widespread change throughout multiple cortical association and projection tracts. Importantly, within individuals, intervention-driven effects are tightly coupled across this collection of tracts. Further, tissue properties and reading skills change in concert: An individual’s time-course of white matter changes tracks their time-course of changes in reading skill. This suggests that the white matter rapidly adapts to the changing environmental demands posed by the intervention. The extent of plasticity in the white matter has important implications for the interpretation of correlations between white matter tissue properties and academic skills: As cross-sectional correlations change week-to-week, correlations measured at any single time point offer an incomplete, and potentially misleading, view of the underlying relationships between anatomy, behavior and experience.

Intervention leads to rapid changes that are distributed across cortical association and projection tracts, including, but not limited to, the left arcuate and left inferior longitudinal fasciculus. These tracts connect distinct components of the reading circuitry, and are generally considered to support separable aspects of reading. For example, the AF has been linked specifically to phonological awareness^8,35^, while the ILF, which projects to the ‘visual word form area’ (VWFA)^47^, may be especially involved in visual word recognition. Typically, over years of development, growth rates for these two tracts are independent from each other^38^. We therefore hypothesized that the learning process might differentially affect tissue properties within these tracts. Further, given the diversity of behavioral profiles seen in people with dyslexia, subjects could show differing spatial profiles of change. For example, a subject with strong intervention-driven effects within the AF might show smaller effects within the ILF, while another subject might show the opposite pattern. However, our results support an alternative view. Longitudinal changes in the AF and ILF are tightly coupled within subjects and also correlated to changes in many other white matter tracts, suggesting that these effects arise from a common biological mechanism operating over a large anatomical scale.

Typically, dMRI studies of the white matter seek to identify a single critical structure that is related to a specific cognitive skill. Our measurements offer a different view on white matter plasticity and learning: Anatomically widespread effects may be a hallmark of rapid, short-term plasticity associated with intensive training of reading skills. Since reading depends on the coordination of a large cortical network, training of reading skills may prompt particularly widespread effects across the white matter. Functional changes measured with fMRI after reading training appear to be widespread^64^, affecting multiple sites within the cortical and sub-cortical reading network. However, a relatively small and focal change in anatomy could theoretically produce widespread functional changes, and therefore these effects need not be accompanied by large-scale anatomical remodeling. Indeed, a small number of past studies in human subjects have reported focal changes in white matter after training of reading skills^32,65^, but past work has not employed the intensive training paradigm used here (see also^33^). Alternatively, the widespread effects may reflect general mechanisms of learning during an intensive educational experience, and therefore may not be specific to the curriculum of this reading intervention.

It is important to note that the tracts identified in this analysis, including the left hemisphere ILF, SLF, and AF, carry signals that are relevant for a number of cognitive functions^66^, not only reading^67–71^. Interestingly, individual differences in plasticity within the left AF have recently been linked to individual gains in math skills following math intervention^72^, even though the left AF is conventionally associated with language related skills. It should be noted, however, that in^72^, math skills training did not produce a significant change in the arcuate at a group level, and therefore the previous set of findings differ from ours. Given the relatively coarse (mm) scale of dMRI, it is possible that distinct types of intervention (e.g., training in reading versus math skills) affect distinct subpopulations of fibers with distinct cortical terminations and functional roles. However, an alternative interpretation also emerges from the current study: Intensive training may lead to plasticity within regions that are not necessarily critical for performing the trained task, and thus intervention-driven effects in the left AF might reflect general mechanisms that are common to learning both reading and math. Despite the lack of a group-level intervention effect in the left arcuate in^72^, it remains possible that a sufficiently intense math intervention might prompt changes not only within the left arcuate, but within many of the same tracts identified here. Indeed, our effects may reflect the intensity and quality of the learning environment, rather than the specific trained skills. Moreover, since it would not be feasible to enroll skilled readers in a highly intensive reading intervention program, it is unclear whether the observed effects are specific to individuals with reading difficulties. Future work examining the generalizability of these effects in other domains, such as math, would allow an examination of general learning effects in a broader population and should help clarify the role of domain specific deficits.

What biological mechanism might underlie the observed effects? Changes in the diffusion signal can arise from multiple sources, including use-related swelling and branching of glial cells^30,31,73^, changes in vasculature, myelination of unmyelinated axons, myelin remodeling, and/or growth of new myelinating oligodendrocytes (reviewed in^74^). Oligodendrocyte precursor cells are present throughout the white matter, and large-scale proliferation of oligodendrocytes has been shown in mice within hours of optogenetically-stimulated activity in adult motor cortex^23^. Mature oligodendrocytes, in turn, participate in myelin maintenance and remodeling throughout the lifespan. Thus, a particularly intriguing possibility is that an initial pattern of widespread change in diffusion properties reported here might reflect rapid growth of myelinating oligodendrocytes, of which only a fraction will ultimately contribute to new myelin sheaths within focal, task relevant regions. In that case, it should be possible to differentiate diffusion signal changes related to rapid growth of oligodendrocytes from signal changes related to longer-term changes in myelin after the training period has ended. In particular, we might expect a rapid initial large decrease in MD, since diffusion would be hindered by new oligodendrocytes. Subsequent changes in myelin might emerge as relatively smaller, persistent changes in other quantitative MRI parameters ^62,75–77^.

Intervention-driven changes in white matter do not follow the trajectory predicted by a normalization account, in which remediation of reading difficulties could be expected to eliminate differences in the white matter between children with dyslexia and typical readers. Instead, we find the opposite: Short-term learning-induced changes are large relative to baseline group differences, and they deviate from the normalization prediction, as shown in Figure 3. Previously reported group differences may therefore be driven in large part by environmental differences between groups, since systematic difference in the environment (e.g., differences in the quality or intensity of recent educational experiences for dyslexic versus control subjects) could be expected to exert a large influence on diffusion measurements, and to potentially counteract or change pre-existing anatomy-behavior relationships. This offers an explanation for why some studies find a positive correlation between FA and reading skills^14,19^, while other studies find a negative correlation between FA and reading skills^8,78^ within the exact same tracts.

In contrast to the widespread changes described above, we find that the posterior callosal connections are remarkably stable over the course of intervention, and also show stable correlations with reading skills. Although we interpret this null result cautiously, one possibility is that differences in MD within posterior callosal connections reflect relatively stable anatomical variation, which predicts reading skill, but does not change during shortterm, intensive training. Indeed, the structure of the posterior corpus callosum differs in both children and adults with dyslexia, and the positive correlation between diffusion properties in this pathway and reading skills has been reported by many other studies^3,4,79–81^. These connections are known to mature relatively early; therefore, the subjects in our study may already be outside the sensitive period in which experience shapes these connections. In that case, training at an earlier age might prompt changes in the CC alongside acquisition of reading skills.

In summary, our results show that altering a child’s educational environment through a targeted intervention program induces rapid, large-scale changes in white matter tissue properties. We observe changes in both MD and FA that occur over the timescale of weeks, that track changes in an individual’s reading skills, and are tightly coupled across tracts connecting distinct parts of the neural circuitry for reading.

## Methods

### Participants

A total of 93 behavioral and MRI sessions were conducted with a group of 24 children (11 female), ranging in age from 7 to 12 years, who participated in an intensive summer reading intervention program. Members of the intervention group were recruited based on parent report of reading difficulties and/or a clinical diagnosis of dyslexia. An additional 52 behavioral and MRI sessions were conducted with 19 participants, who were matched for age but not reading level. These subjects were recruited as a control group to assess the stability of our measurements over the repeated sessions. Control subjects participated in the same experimental sessions, but did not receive the reading intervention. Ten of these subjects had typical reading skills (4 female), defined as a score of 85 or greater on the Woodcock Johnson Basic Reading composite and the TOWRE Index. Nine had reading difficulties (3 female), defined as a score below 85 on either the Woodcock Johnson Basic Reading composite or the TOWRE Index. Reading assessments were carried out at the start of the intervention period to confirm parent reports and establish a baseline for subsequent estimates of growth in reading skill. Demographics and initial test scores are summarized in **Table 2**.

All participants were native English speakers with normal or corrected-to-normal vision and no history of neurological damage or psychiatric disorder. We obtained written consent from parents, and verbal assent from all child participants. All procedures, including recruitment, consent, and testing, followed the guidelines of the University of Washington Human Subjects Division and were reviewed and approved by the UW Institutional Review Board.

**Table 2.** Subject demographics and pre-intervention test scores. Age and pre-intervention Woodcock Johnson Basic Reading Skills, Reading Fluency, and Test of Word Reading Efficiency (TOWRE) standard scores are given for the intervention and control groups. Intervention and control groups were matched in age, but the intervention group had significantly lower reading scores.

| **Subject Group** | **Age in years**<br>**(mean/std.dev.)** | **Basic Reading**<br>**(mean/std.dev.)** | **Reading**<br>**Fluency**<br>**(mean/std.dev.)** | **TOWRE**<br>**Composite**<br>**(mean/std.dev.)** |
| --- | --- | --- | --- | --- |
| Intervention | 9.71/1.80 | 79.45/16.27 | 72.05/ 21.065 | 72.25/14.48 |
| Control | 9.95/1.36 | 95.17/16.87 | 91.17/18.63 | 87.44/18.18 |
| Intervention vs.<br>Control | $t(36) = -0.47,$<br>$p = 0.64$ | $t(36) = -2.92,$<br>$p = 0.0060$ | $t(36) = -2.95,$<br>$p = 0.0056$ | $t(36) = -2.86$<br>$p = 0.0069$ |

### Reading intervention

Intervention subjects were enrolled in 8 weeks of the *Seeing Stars: Symbol Imagery for Fluency, Orthography, Sight Words, and Spelling*^82^ program at three different *Lindamood-Bell Learning Centers* in the Seattle area. The intervention program consists of directed, one-on-one training in phonological and orthographic processing skills, lasting four hours each day, five days a week. The curriculum uses an incremental approach, building from letters and syllables to words and connected texts, emphasizing phonological decoding skills as a foundation for spelling and comprehension. A hallmark of this intervention program is the intensity of the training protocol (4 hours a day, 5 days a week) and the personalized approach that comes with one-on-one instruction.

### Experimental Sessions

Subjects participated in four experimental sessions separated by roughly 2.5-week intervals. For the intervention group, sessions were scheduled to occur before the intervention (baseline), after 2.5 weeks of intervention, after 5 weeks of intervention, and at the end of the 8-week intervention period. For the control group, sessions followed the same schedule while the subjects attended school as usual. This allowed us to control for changes that would occur due to typical development and learning during the school year. Twenty-one intervention subjects completed all four experimental sessions; 3 subjects completed only 3 sessions, which fell at the start, middle and end of the intervention. In the control group, 7 subjects completed all 4 sessions; 12 subjects completed at least 3 sessions; 14 subjects completed at least 2 sessions; 19 subjects completed at least one session.

In addition to MRI measurements, described in greater detail below, we administered a battery of behavioral tests in each experimental session. These included sub-tests from the Wechsler Abbreviated Scales of Intelligence (WASI), Comprehensive Test of Phonological Processing (CTOPP-2), Test of Word Reading Efficiency (TOWRE-2) and the Woodcock Johnson IV Tests of Achievement (WJ-IV). Rather than analyzing each subtest individually, we created a general reading skills index by conducting a principal component analysis on subtests from the latter two batteries (TOWRE and WJ-IV) and taking scores from the first principal component, which accounted for 83.76% of the total variance in reading performance (**Supplementary Figure 1**). We used this measure for all subsequent analysis in order to avoid issues that arise from multiple comparisons, and to increase the reliability of our reading skill index. Our Reading Skills index was highly correlated with both the WJ-BRS composite (*r*(97) = 0.95, *p* < 0.001) and the TOWRE composite (*r*(97) = 0.96, *p* < 0.001).

### MRI Acquisition and Processing

All imaging data were acquired with a 3T Phillips Achieva scanner (Philips, Eindhoven, Netherlands) at the University of Washington Diagnostic Imaging Sciences Center (DISC) using a 32-channel head coil. An inflatable cap was used to minimize head motion, and participants were continuously monitored through a closed circuit camera system. Prior to the first MRI session, all subjects completed a session in an MRI simulator, which helped them to practice holding still, with experimenter feedback. This practice session also allowed subjects to experience the noise and confinement of the scanner prior to the actual imaging sessions, to help them feel comfortable and relaxed during data collection.

Diffusion-weighted magnetic resonance imaging (dMRI) data were acquired with isotropic 2.0mm^3^ spatial resolution and full brain coverage. Each session consisted of 2 DWI scans, one with 32 non-collinear directions (b-value = 800 s/mm^2^), and a second with 64 non-collinear directions (b-value = 2,000 s/mm^2^). The gradient directions were optimized to provide uniform coverage^83^. Each of the DWI scans also contained 4 volumes without diffusion weighting (b-value = 0). In addition, we collected one scan with 6 non-diffusion-weighted volumes and a reversed phase encoding direction (posterior-anterior) to correct for EPI distortions due to inhomogeneities in the magnetic field. Distortion correction was performed using FSL’s *topup* tool^84^. Additional pre-processing was carried out using tools in FSL for motion and eddy current correction^85^, and diffusion metrics were fit using the diffusion kurtosis model^86^ as implemented in DIPY^87^. Data were manually checked for imaging artifacts and excessive dropped volumes. Given that subject motion can be especially problematic for the interpretation of group differences in DWI data^88^, data sets with mean slice-by-slice RMS displacement > 0.7mm were excluded from all further analyses. Datasets in which more than 10% of volumes were either dropped or contained visible artifact were also excluded from further analysis. In total, these criteria removed 13 out of 93 total intervention datasets, and 3 out of 52 control datasets.

To further quantify potential effects of motion, we tested for differences in motion across sessions and subject groups (intervention vs. control; see **Supplementary Figure 3**), after excluding datasets based on the criteria listed above. We observed no difference in motion as a function of session (*F*(3,121) = 0.090, *p* = 0.97) or group (*F*(1,121) = 2.54, *p* = 0.11), and no group-by-session interaction (*F*(3,121) = 0.30, *p* = 0.83). Thus, we do not attribute the between-session changes in white matter within the intervention group to systematic differences in motion. Further, including motion as a covariate in our analysis did not change our results, as described below.

### White Matter Tract Identification

Fiber tracts were identified for each subject using the Automated Fiber Quantification (AFQ) software package^38^, after initial generation of a whole-brain connectome using probabilistic tractography (MRtrix 3.0)^89^. Fiber tracking was carried out on an aligned, distortion corrected, concatenated dataset including all four of the 64-direction (b-value = 2,000 s/mm^2^) datasets collected across sessions for each subject. This allowed us to ensure that estimates of diffusivity and diffusion anisotropy across session were mapped to the same anatomical location for each subject, since slight differences in diffusion properties over the course of intervention can influence the region of interest that is identified by the tractography algorithm. We also replicated our main results using tractography derived separately for each session and subject (see **Supplementary Figure 4**).

We focused our initial analysis on 3 tracts that are thought to connect the core reading circuitry ^37,38^: the left arcuate fasciculus (AF), left inferior longitudinal fasciculus (ILF), and posterior callosal connections (CC). Subsequent analysis included 13 additional tracts: Left and right thalamic radiations, left and right corticospinal tracts, anterior callosal connections, left and right inferior frontal occipital fasciculus (IFOF), right ILF, left and right superior longitudinal fasciculus (SLF), left and right uncinate, and right AF.

### Quantifying White Matter Tissue Properties

To detect intervention-driven changes in the white matter, we fit the diffusion kurtosis model^86^ as implemented in DIPY^87^ to the diffusion data collected in each session. The diffusion kurtosis model is an extension of the diffusion tensor model that accounts for the non-Gaussian behavior of water in heterogeneous tissue containing multiple barriers to diffusion (cell membranes, myelin sheaths, etc.). After model fitting, diffusion metrics were projected onto the segmented fiber tracts generated by AFQ. Selected tracts were sampled into 100 evenly spaced nodes, spanning termination points at the gray-white matter boundary, and then diffusion properties (mean, radial, and axial diffusivity (MD, RD, AD) and fractional anisotropy (FA)) were mapped onto each tract to create a “Tract Profile”.

### Code and Data Availability

All code and data required to reproduce reported findings is available at **[URL available upon publication]**.

### Statistical Analysis

Data analysis was carried out using software written in MATLAB. To assess change over the course of intervention, we first averaged the middle 60% of each tract to create a single estimate of diffusion properties for each subject and tract. We selected the middle portion to eliminate the influence of crossing fibers near cortical terminations, and to avoid potential partial volume effects at the white matter / gray matter border. Mean tract values were then entered into a linear mixed effects model, with fixed effects of intervention time (either hours of training, or session entered as a categorical variable) and a random effect of subject. We modeled the relationship between white-matter properties and behavior in a similar fashion, predicting Reading Skill from mean tract values and session, with subjects treated as a random effect.

We further examined the time-course of change in white matter and reading skills by (1) performing a cross-correlation analysis on individual longitudinal trajectories and (2) calculating individual linear growth rates, which allowed us to directly model relationships between behavioral and white-matter growth rates across subjects.

Finally, to examine the anatomical specificity of intervention-driven changes, we fit a mixed linear model to the growth trajectories of a large collection of white matter tracts. We then performed hierarchical clustering on the correlations between linear growth-rates, using a complete-linkage clustering algorithm implemented in MATLAB, to test for correlated growth trajectories across a large collection of cortical association tracts.

## SUPPLEMENTARY FIGURES AND TABLES

**Figure S1.**
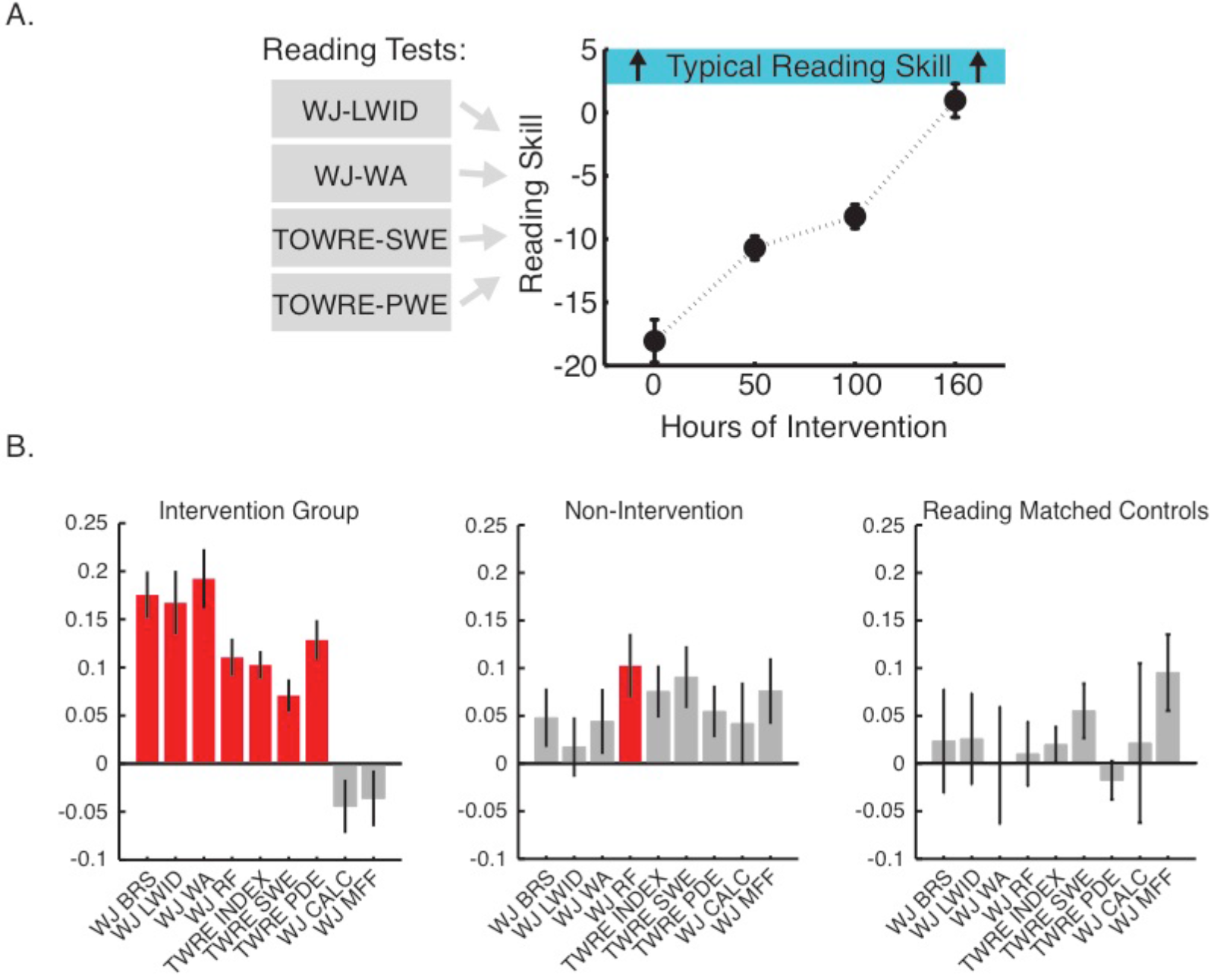
Intervention driven growth in reading performance. (A) We created a single summary index of reading skills based on conducting principal component analysis of the Woodcock Johnson and Test of Word Reading Efficiency standard scores (see Methods). Intervention driven change in this Reading Skill composite is plotted as a function of intervention hours and shows highly significant change (linear mixed effects model, fixed effect of intervention hours and random effect of subject, p<10^−9^). (B) Linear growth in each of the standardized reading measures comprising the Reading Skill composite. In the intervention group, each of the reading subtests grew significantly during the intervention. For the full sample of non-intervention control subjects, we found a significant increase in performance in timed measures, reflecting practice with the tests. However, this practice-related improvement was only present for the children with good reading skills, and there was no change in any of the reading measures in the reading-skill matched control group.

**Figure S2.**
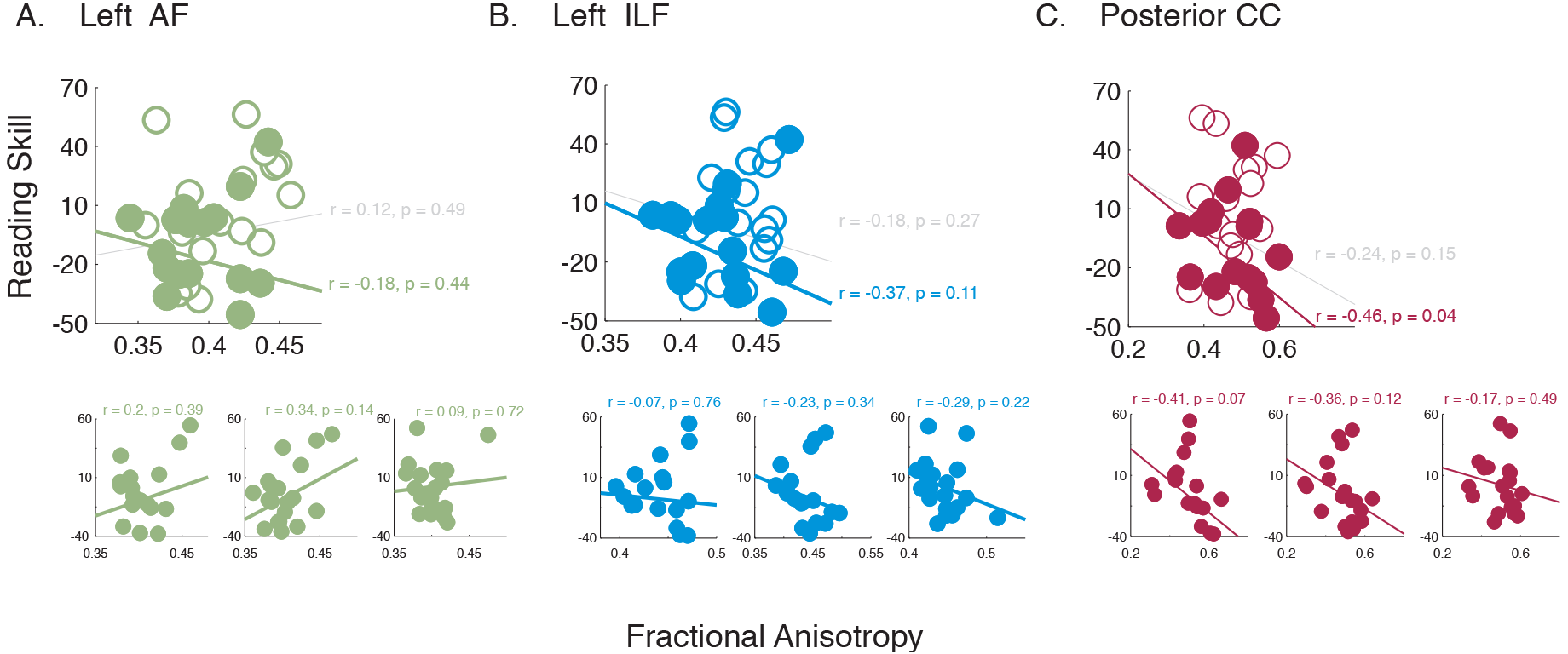
Tract properties vary as a function of pre-intervention reading skill. (A-C) Tract average FA is plotted as a function of pre-intervention (Session 1) reading skill. Best-fit lines plotted in gray give estimates for the full data set, while colored lines show fits for the intervention subjects, alone. Correlations for the intervention subjects are given in colored text below the value estimated for the full data set (in gray). Insets, below, show the cross-sectional correlation for Sessions 2-4 (left to right), during the intervention. Correlation values are reported for the intervention subjects, to highlight changes in the anatomy-behavior correlations that are specific to the intervention group.

**Figure S3.**
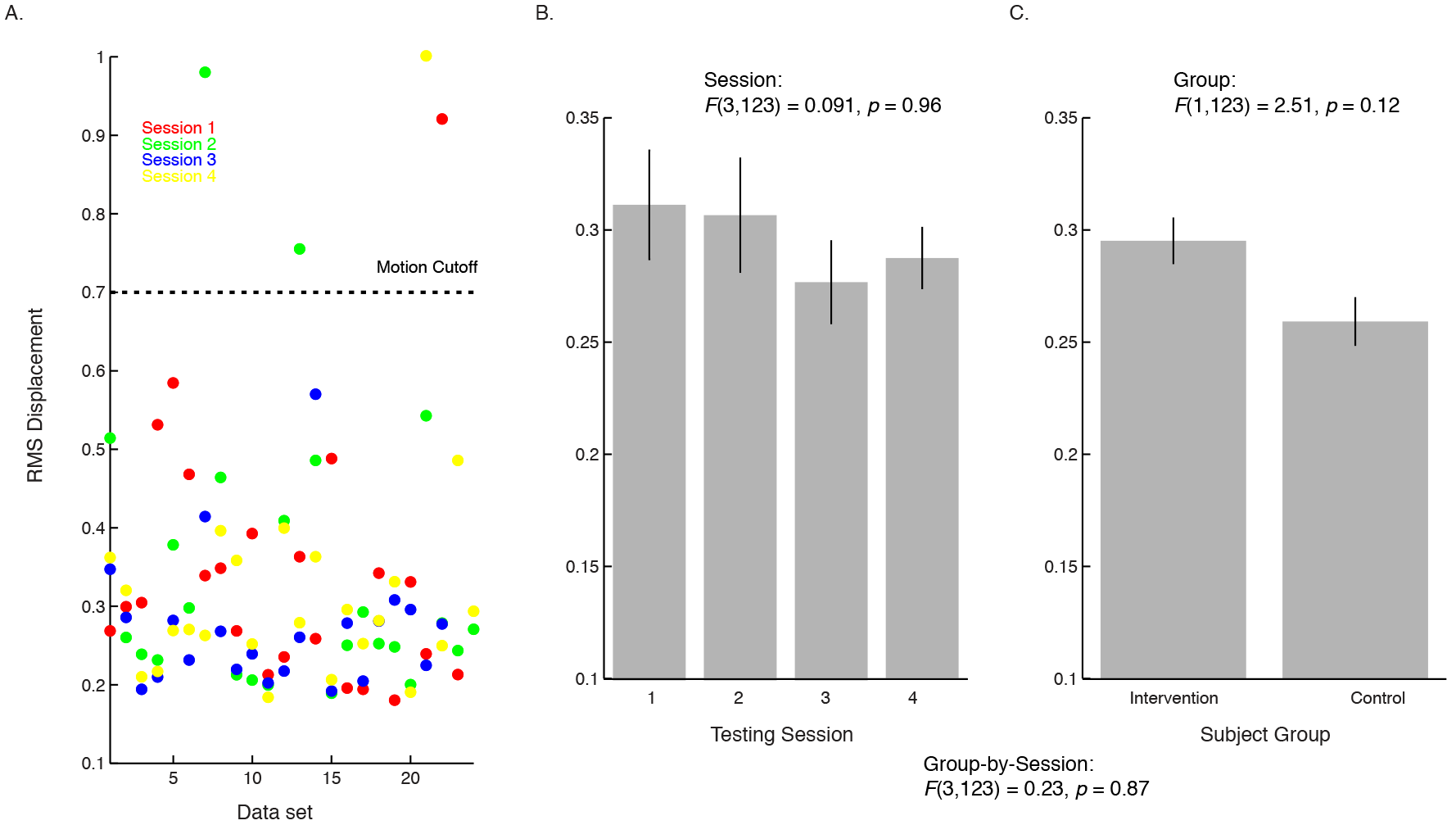
Analysis of subject motion. (A) Mean RMS displacement plotted for each intervention subject and session. Sessions are color coded (sessions 1-4 in red, blue, green, and yellow). Data sets included in subsequent analysis met the following criteria: (1) mean slice-by-slice RMS displacement < 0.7mm, (2) < 10% of volumes dropped or contained visible artifact. In total, we removed 9 of 93 total intervention datasets, and 3 of 52 control datasets. (B) Mean RMS motion is plotted for each session. Error bars represent standard error of the mean. Head motion did not differ across intervention sessions 1-4. (C) We found no group-by-session interaction in head motion, suggesting that changes in diffusivity within training, in the intervention group, could not be attributed to motion.

**Figure S4.**
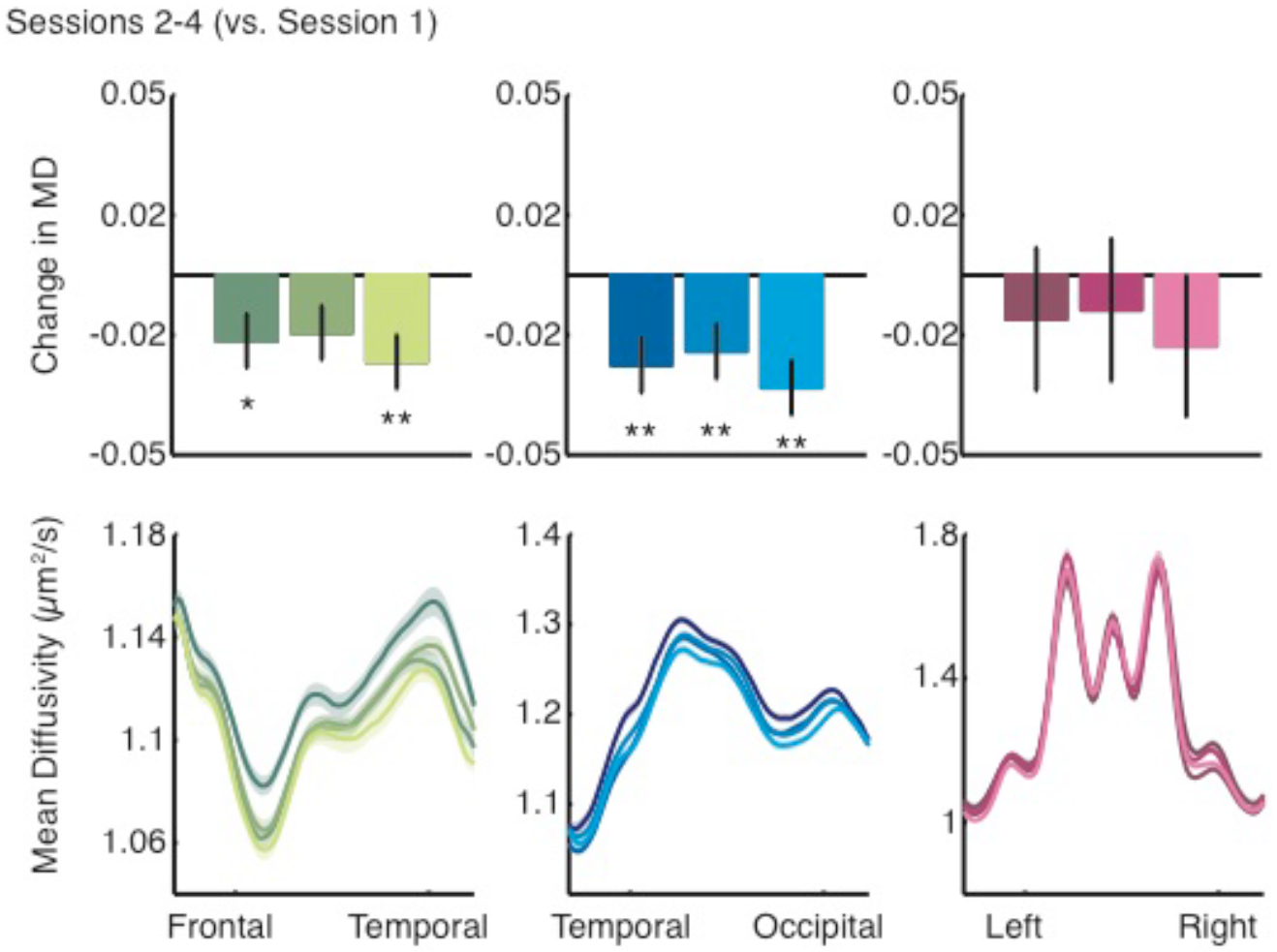
Changes in MD plotted on single session tractography for the intervention group. Intervention-driven changes do not depend on whether fiber tracts are identified independently for each session or in the multi-session concatenated data. (A-B) Values are selected based on tractography carried out on same-session data. Note that the regions of interest identified by the tractography algorithm will differ slightly between sessions. (A) Model predicted change in MD for each session (relative to baseline). Asterisks indicate significant decrease in MD for each session relative to the pre-intervention measurement (Session 1) at a Bonferroni corrected p < 0.05. (B) Tract profiles showing average mean diffusivity (MD) across subjects, measured at four time-points: pre-intervention (Session 1), after 2.5 weeks of intervention (Session 2), after 5 weeks of intervention (Session 3), and after 8 weeks of intervention (Session 4). Shaded error bars give ±1 standard error of the mean. Color values indicate session, ranging from darkest (Session 1) to brightest (Session 4) for each tract. Both the arcuate fasciculus (AF) and inferior longitudinal fasciculus (ILF) show a systematic decrease in MD over the course of intervention.

**Figure S5.**
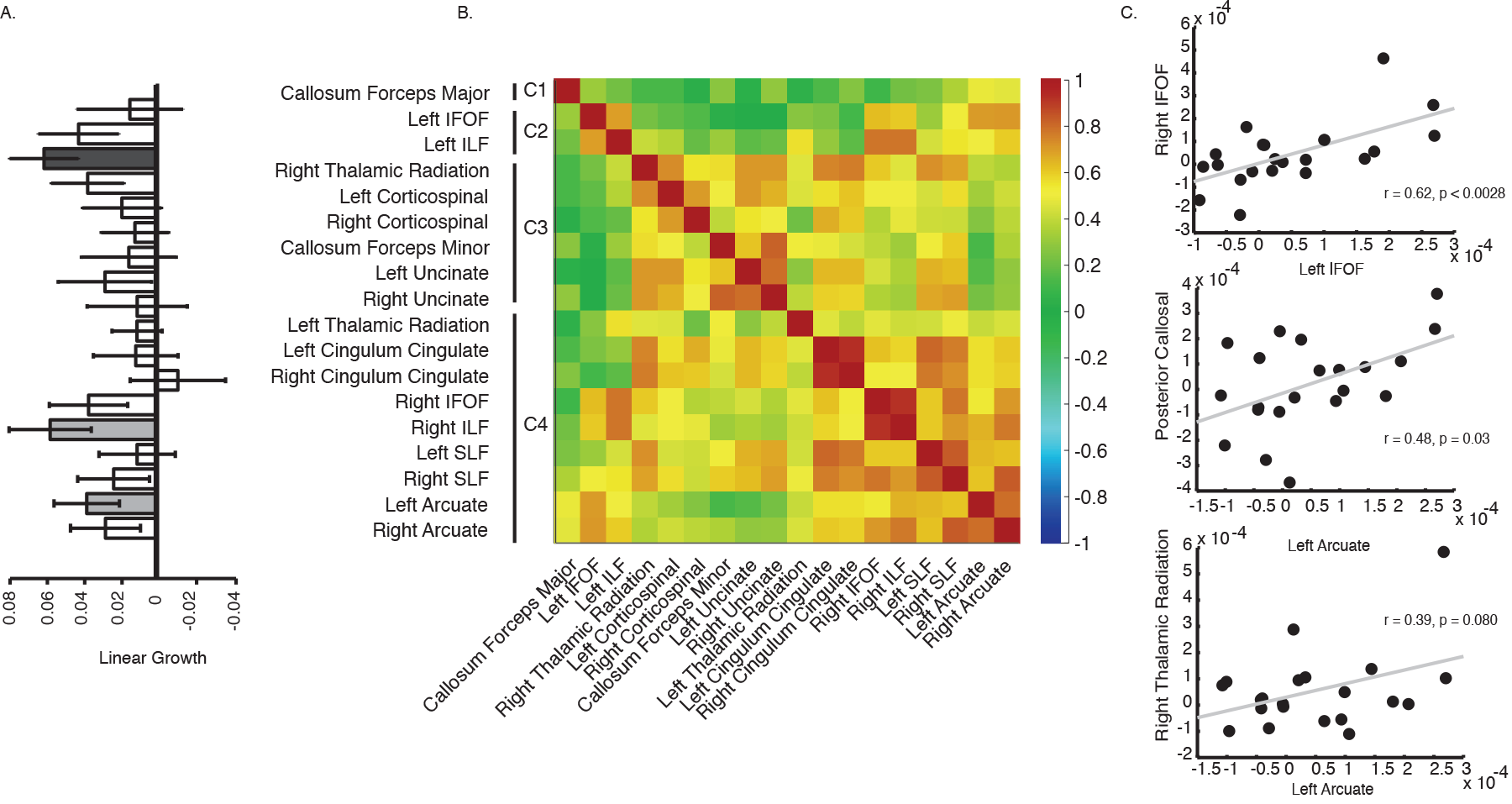
Reading intervention causes distributed changes in the white matter. (A) Change in FA as a function of intervention time (in hours) for 18 tracts. Tracts showing significant change (Bonferroni corrected p < 0.05) are indicated as dark gray filled bars. Tracts showing change at an uncorrected threshold (p < 0.05) are indicated as light gray filled bars. (B) Hierarchical clustering based on the correlations between linear growth rates. The heat map represents correlations between linear growth rates for pairs of tracts. The matrix is sorted according to hierarchical clustering of these correlations. (C) Scatter plots of individual growth rates for three pairs of tracts: left vs. right IFOF, AF vs. CC, and AF vs. right thalamic radiation.

**Figure S6.**
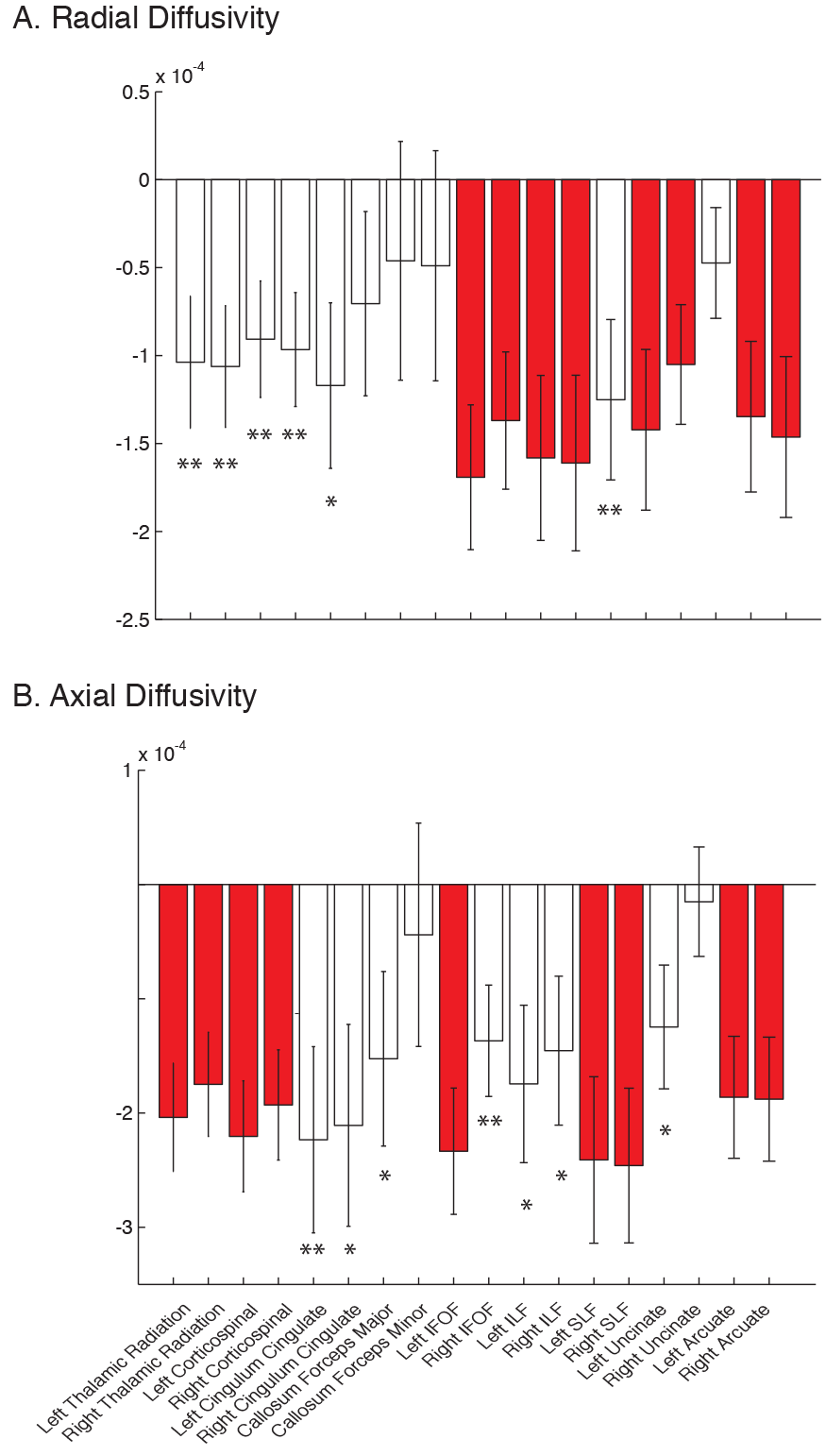
Change in additional diffusion as a function of intervention time (in hours) for individual tracts. Tracts showing significant change (Bonferroni corrected p < 0.05) are highlighted in red. Tracts with significant change before correction are indicated with a single (uncorrected p < 0.05) or double (uncorrected p < 0.01) asterisk.

**Table S1.**
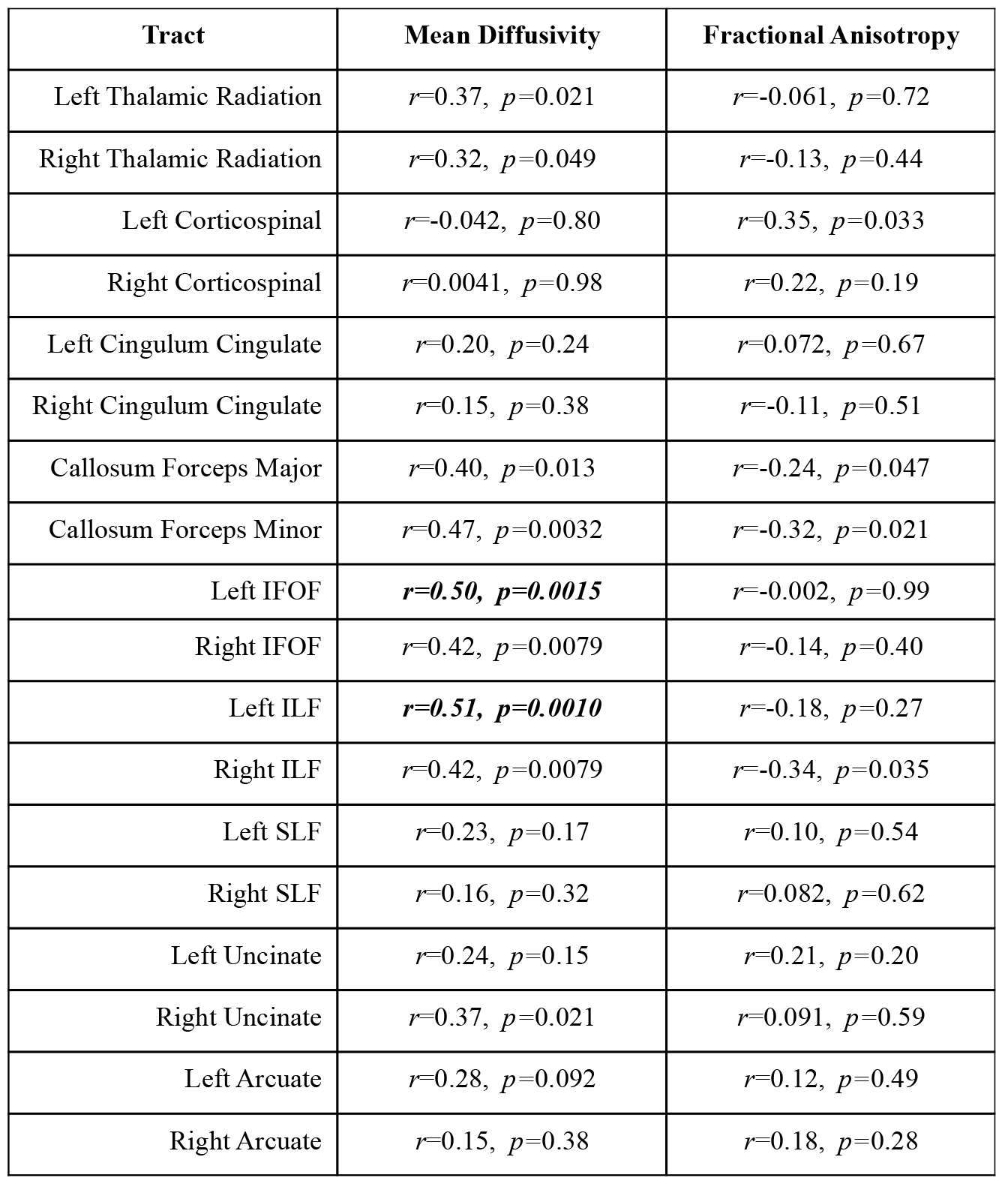
Tract properties and reading skills are correlated at baseline (pre-intervention). Correlations with pre-intervention MD and FA are given for all tracts included in an exploratory analysis. Statistics were calculated using the full (Intervention and Control) sample of baseline (pre-intervention) Reading Skill scores. Tracts showing significant correlation at a conservative threshold (Bonferroni corrected *p* < 0.05) are highlighted in bold italic.

**Table S2.** Behavioral effects. Growth in reading skill was specific to the intervention group. Results for the full control sample (n = 19), including 10 skilled reading control subjects. In this group, performance improved with repeated testing for the timed measures (TOWRE and Reading Fluency). In all control subjects, untimed measures (WJ Basic Reading) were stable, showing no change over 8 weeks.

| Reading Measure | Group | Time | Interaction |
| --- | --- | --- | --- |
| Reading Skill (PCA) | $F(1,124) = 8.55,$<br>$p = 0.0041$ | $F(1,124) = 5.34,$<br>$p = 0.022$ | $F(1,124) = 9.38,$<br>$p = 0.0027$ |
| WJ-BRS | $F(1,124) = 9.59,$<br>$p = 0.0024$ | $F(1,124) = 1.86,$<br>$p = 0.17$ | $F(1,124) = 8.64,$<br>$p = 0.0039$ |
| TOWRE Index | $F(1,124) = 6.52,$<br>$p = 0.012$ | $F(1,124) = 9.28,$<br>$p = 0.0028$ | $F(1,124) = 0.85,$<br>$p = 0.36$ |
| WJ-CALC | $F(1,98) = 8.79,$<br>$p = 0.0038$ | $F(1,98) = 0.92,$<br>$p = 0.34$ | $F(1,98) = 2.78,$<br>$p = 0.099$ |

**Table S3.**
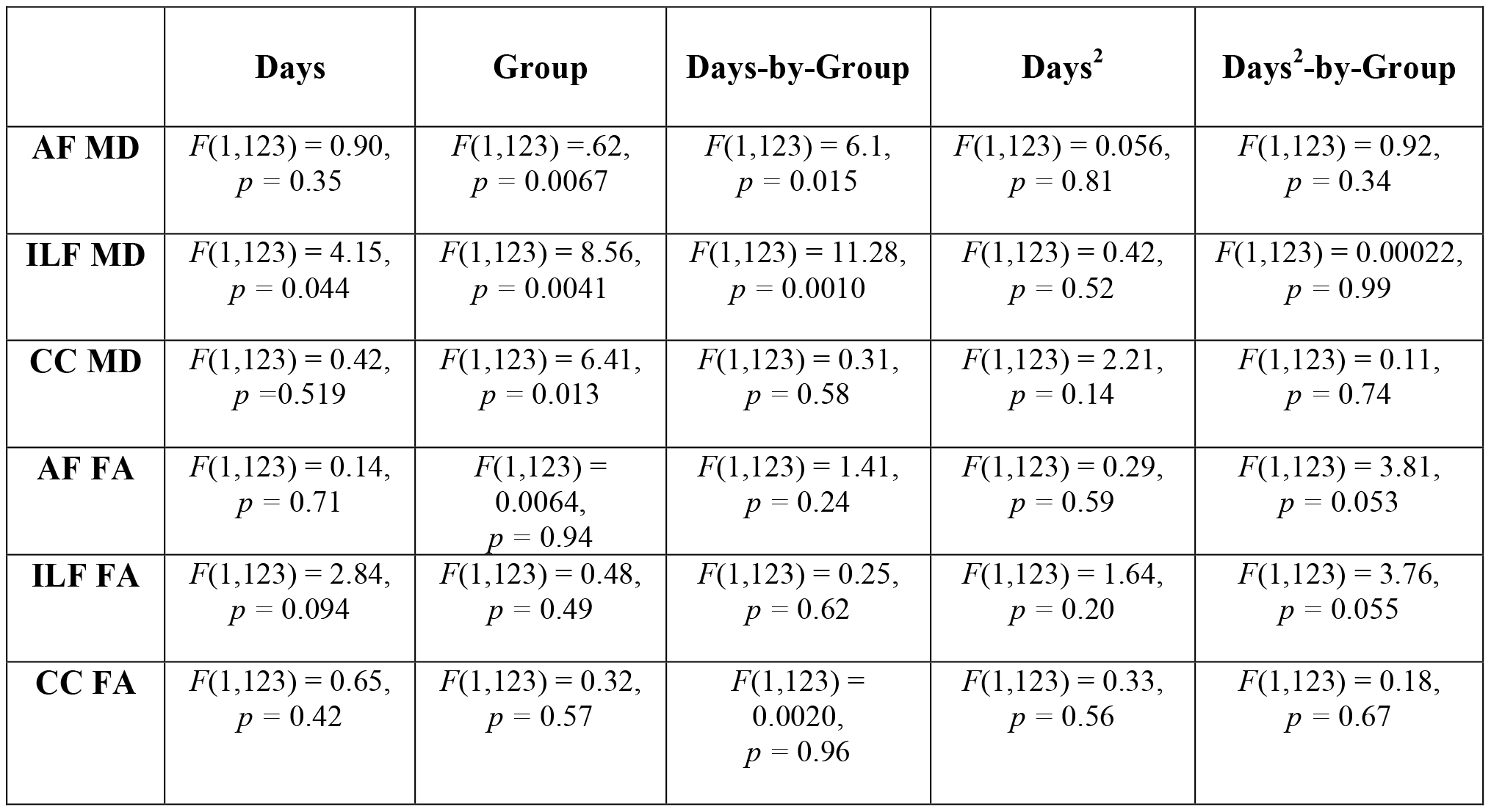
Linear and quadratic effects in the white matter. Results for a linear mixed effects model predicting white matter properties (mean diffusivity, MD, and fractional anisotropy, FA) for the left arcuate (AF), inferior longitudinal fasciculus (ILF) and posterior callosal connections (CC) as a function of intervention time (in days) and subject group (intervention vs. non-intervention control),

**Table S4.** Variance in white matter and reading skills is well matched across sessions. F-statistics in each cell represent the ratio of variance across time points for each white matter tract, and the Reading Skill composite, calculated with the larger of the 2 variances in the numerator. Statistics were computed using the 20 intervention subjects with the full set of 4 MRI data points.

|  | **Reading Composite** | **Posterior CC** | **Left ILF** | **Left Arcuate** |
| --- | --- | --- | --- | --- |
| **Session 1 vs. 2** | $F(19,19) = 1.088$,<br>$p > 0.05$ | $F(19,19) = 1.17$,<br>$p > 0.05$ | $F(19,19) = 1.045$,<br>$p > 0.05$ | $F(19,19) = 1.16$,<br>$p > 0.05$ |
| **Session 1 vs. 3** | $F(19,19) = 1.24$,<br>$p > 0.05$ | $F(19,19) = 1.30$,<br>$p > 0.05$ | $F(19,19) = 1.25$,<br>$p > 0.05$ | $F(19,19) = 1.075$,<br>$p > 0.05$ |
| **Session 1 vs. 4** | $F(19,19) = 1.49$,<br>$p > 0.05$ | $F(19,19) = 1.70$,<br>$p > 0.05$ | $F(19,19) = 1.98$,<br>$p > 0.05$ | $F(19,19) = 2.27$,<br>$p > 0.01$ |

**Table S5.** Baseline normalized analysis of learning trajectories. Results of a mixed effects model predicting mean diffusivity, relative to baseline, in the left inferior longitudinal fasciculus (ILF), and the left arcuate (AF) from reading skill throughout the intervention, and mean diffusivity, relative to baseline, in the left inferior longitudinal fasciculus (ILF) as a function of mean diffusivity, relative to baseline, in the left arcuate (AF).

|  | Effect in the Intervention Group |
| --- | --- |
| **ILF vs. Reading Skill** | $F(1,68) = 4.46, p = 0.038$ |
| **AF vs. Reading Skill** | $F(1,68) = 3.98, p = 0.050$ |
| **ILF vs. AF** | $F(1,69) = 335.42, p > 10^{-27}$ |

**Table S6.** Non-Intervention MD and FA Versus Reading Skill Over Time. Cells show p-values based on a mixed linear model predicting session-to-session changes in Reading Skill composite scores from in mean diffusivity (MD) and fractional anisotropy (FA) in the non-intervention Control subjects. Pearson correlations between mean-centered MD/FA and mean-centered reading score are provided as an index of effect size. Consistent with the stability of both reading scores and diffusion properties in the control group, no tracts change in relation to Reading Skill in the Control group.

| Tract | Mean Diffusivity | Fractional Anisotropy |
| --- | --- | --- |
| Left Thalamic Radiation | $r=-0.035, p=0.81$ | $r=0.0016, p=0.99$ |
| Right Thalamic Radiation | $r=0.079, p=0.58$ | $r=-0.17, p=0.22$ |
| Left Corticospinal | $r=-0.052, p=0.72$ | $r=0.017, p=0.91$ |
| Right Corticospinal | $r=0.062, p=0.67$ | $r=-0.025, p=0.86$ |
| Left Cingulum Cingulate | $r=-0.042, p=0.77$ | $r=0.011, p=0.94$ |
| Right Cingulum Cingulate | $r=0.13, p=0.36$ | $r=-0.027, p=0.85$ |
| Callosum Forceps Major | $r=-0.15, p=0.29$ | $r=0.15, p=0.31$ |
| Callosum Forceps Minor | $r=0.030, p=0.84$ | $r=-0.049, p=0.73$ |
| Left IFOF | $r=-0.012, p=0.93$ | $r=0.081, p=0.081$ |
| Right IFOF | $r=0.17, p=0.23$ | $r=-0.023, p=0.87$ |
| Left ILF | $r=0.070, p=0.62$ | $r=0.021, p=0.88$ |
| Right ILF | $r=0.0013, p=0.99$ | $r=-0.057, p=0.69$ |
| Left SLF | $r=-0.080, p=0.58$ | $r=-0.045, p=0.76$ |
| Right SLF | $r=0.053, p=0.71$ | $r=-0.048, p=0.74$ |
| Left Uncinate | $r=-0.019, p=0.90$ | $r=0.0050, p=0.97$ |
| Right Uncinate | $r=0.099, p=0.49$ | $r=-0.16, p=0.25$ |
| Left Arcuate | $r=-0.046, p=0.75$ | $r=-0.059, p=0.68$ |
| Right Arcuate | $r=0.059, p=0.68$ | $r=-0.13, p=0.38$ |

**Table S7.**
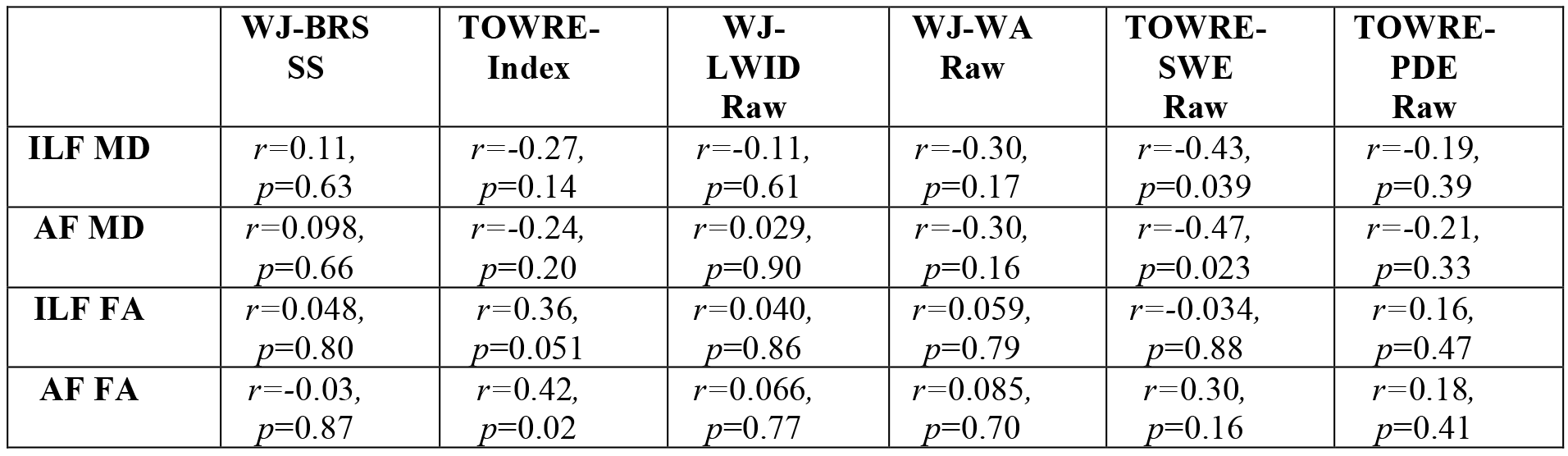
Session 2 vs. Session 1 Difference Scores. Each cell gives the Session 2 vs. Session 1 difference score for the specified tract (left arcuate, AF, or left ILF) and parameter (mean diffusivity, MD, or fractional anisotropy, FA) and reading measure (Woodcock-Johnson or TOWRE subtests, from left to right: WJ Basic Reading Standard Score, TOWRE Index, WJ Letter-Word ID, WJ Word Attack, TOWRE Sight Word Efficiency, TOWRE Phonemic Decoding Efficiency).

**Table S8.** FA Change Versus Reading Skill Change during the intervention. Cells show p-values based on a mixed linear model predicting session-to-session changes Reading Skill composite score, Woodcock-Johnson Basic Reading scores, and TOWRE index scores, and Reading Fluency from changes in fractional anisotropy (FA) at each time point during the intervention. Pearson correlations between mean-centered FA and mean-centered reading score are provided as an index of effect size.

| Tract (FA) | Reading Skill Composite | Basic Reading Score | TOWRE Index | Reading Fluency |
| --- | --- | --- | --- | --- |
| Left Thalamic Radiation | $r=0.020, p=0.86$ | $r=-0.061, p=0.59$ | $r=0.095, p=0.40$ | $r=0.031, p=0.78$ |
| Right Thalamic Radiation | $r=0.13, p=0.25$ | $r=0.082, p=0.46$ | $r=0.12, p=0.30$ | $r=0.30, p=0.0076$ |
| Left Corticospinal | $r=0.14, p=0.20$ | $r=0.067, p=0.55$ | $r=0.15, p=0.19$ | $r=0.21, p=0.065$ |
| Right Corticospinal | $r=0.060, p=0.60$ | $r=0.055, p=0.62$ | $r=-0.0020, p=0.99$ | $r=0.16, p=0.14$ |
| Left Cingulum Cingulate | $r=0.045, p=0.69$ | $r=0.029, p=0.80$ | $r=-0.0060, p=0.96$ | $r=0.044, p=0.70$ |
| Right Cingulum Cingulate | $r=-0.093, p=0.41$ | $r=-0.14, p=0.23$ | $r=-0.036, p=0.75$ | $r=-0.0074, p=0.95$ |
| Callosum Forceps Major | $r=0.014, p=0.90$ | $r=-0.055, p=0.62$ | $r=0.15, p=0.19$ | $r=0.12, p=0.30$ |
| Callosum Forceps Minor | $r=-0.038, p=0.74$ | $r=-0.058, p=0.61$ | $r=0.047, p=0.68$ | $r=0.063, p=0.58$ |
| Left IFOF | $r=0.10, p=0.36$ | $r=0.098, p=0.39$ | $r=0.11, p=0.31$ | $r=0.14, p=0.23$ |
| Right IFOF | $r=0.13, p=0.24$ | $r=0.13, p=0.24$ | $r=0.14, p=0.20$ | $r=0.26, p=0.021$ |
| Left ILF | $r=0.20, p=0.08$ | $r=0.13, p=0.25$ | $r=0.26, p=0.019$ | $r=0.19, p=0.093$ |
| Right ILF | $r=0.17, p=0.14$ | $r=0.14, p=0.23$ | $r=0.18, p=0.10$ | $r=0.26, p=0.020$ |
| Left SLF | $r=0.032, p=0.77$ | $r=-0.039, p=0.73$ | $r=0.062, p=0.58$ | $r=0.051, p=0.66$ |
| Right SLF | $r=0.10, p=0.36$ | $r=0.088, p=0.44$ | $r=0.063, p=0.58$ | $r=0.14, p=0.21$ |
| Left Uncinate | $r=0.15, p=0.18$ | $r=0.17, p=0.12$ | $r=0.028, p=0.80$ | $r=0.29, p=0.0096$ |
| Right Uncinate | $r=-0.063, p=0.58$ | $r=-0.020, p=0.86$ | $r=-0.17, p=0.12$ | $r=-0.021, p=0.86$ |
| Left Arcuate | $r=0.078, p=0.49$ | $r=-0.016, p=0.89$ | $r=0.17, p=0.12$ | $r=0.078, p=0.49$ |
| Right Arcuate | $r=0.11, p=0.32$ | $r=0.040, p=0.72$ | $r=0.19, p=0.096$ | $r=0.11, p=0.33$ |

**Table S9:** Individual rates of change in diffusion measures vs. Reading Skill. Each subject’s rate of change for the Reading Skill composite, and for each tract (left arcuate, AF, or left ILF) and parameter (mean diffusivity, MD, or fractional anisotropy, FA), is estimated from the linear fit to intervention hours. All subjects’ rates of change in reading and white matter properties are then used to estimate the correlation between the magnitude of change in reading and white matter.

|  | Correlation between individual rates of change |
| --- | --- |
| **ILF MD** | $r = 0.11, p = 0.65$ |
| **AF MD** | $r = -0.066, p = 0.78$ |
| **ILF FA** | $r = -0.19, p = 0.41$ |
| **AF FA** | $r = -0.16, p = 0.49$ |

**Table S10.** Exploratory Analysis of Intervention-Driven Change. Intervention-driven changes in the white matter are quantified based on the interaction effect (Group-by-Session; linear-mixed effects model with fixed effects of Group (Intervention vs. Control) and Session (1-2), and random effects of subject) in a large collection of white matter tracts. Intervention driven change in MD and FA is reported for each tract. For this exploratory analysis we use a conservative Bonferroni correction, and tracts showing significant change in the Intervention group (Bonferroni corrected *p* < 0.05) are highlighted in bold italic.

| Tract | Mean Diffusivity | Fractional Anisotropy |
| --- | --- | --- |
| Left Thalamic Radiation | $F(1,67)=6.38, p=0.014$ | $F(1,67)=0.44, p=0.51$ |
| Right Thalamic Radiation | **$F(1,67)=11.70, p=0.0011$** | $F(1,67)=1.41, p=0.24$ |
| Left Corticospinal | **$F(1,67)=16.69, p=0.00012$** | $F(1,67)=3.13, p=0.081$ |
| Right Corticospinal | $F(1,67)=8.74, p=0.0043$ | $F(1,67)=0.77, p=0.38$ |
| Left Cingulum Cingulate | $F(1,67)=7.26, p=0.0089$ | $F(1,67)=1.05, p=0.31$ |
| Right Cingulum Cingulate | $F(1,67)=4.30, p=0.042$ | $F(1,67)=0.28, p=0.60$ |
| Callosum Forceps Major | $F(1,67)=0.62, p=0.44$ | $F(1,67)=2.38, p=0.13$ |
| Callosum Forceps Minor | $F(1,67)=3.56, p=0.063$ | $F(1,67)=1.96, p=0.17$ |
| Left IFOF | **$F(1,67)=12.29, p=0.00082$** | $F(1,67)=7.02, p=0.010$ |
| Right IFOF | **$F(1,67)=21.28, p<10^{-4}$** | $F(1,67)=2.96, p=0.090$ |
| Left ILF | $F(1,67)=7.75, p=0.0070$ | $F(1,67)=6.45, p=6.45$ |
| Right ILF | **$F(1,67)=9.68, p=0.0027$** | $F(1,67)=4.38, p=0.040$ |
| Left SLF | $F(1,67)=6.56, p=0.013$ | $F(1,67)=0.11, p=0.74$ |
| Right SLF | **$F(1,67)=9.69, p=0.0027$** | $F(1,67)=0.0048, p=0.95$ |
| Left Uncinate | $F(1,67)=4.88, p=0.031$ | $F(1,67)=4.29, p=0.042$ |
| Right Uncinate | $F(1,67)=3.92, p=0.052$ | $F(1,67)=1.69, p=0.20$ |
| Left Arcuate | $F(1,67)=6.91, p=0.011$ | $F(1,67)=2.85, p=0.096$ |
| Right Arcuate | $F(1,67)=8.95, p=0.0039$ | **$F(1,67)=12.27, p=0.00083$** |

## References

1 Klingberg, T. et al. Microstructure of temporo-parietal white matter as a basis for reading ability: evidence from diffusion tensor magnetic resonance imaging. Neuron 25, 493–500 (2000).

2 Niogi, S. N. & McCandliss, B. D. Left lateralized white matter microstructure accounts for individual differences in reading ability and disability. Neuropsychologia 44, 2178–2188, doi: 10.1016/j.neuropsychologia.2006.01.011 (2006).

3 Dougherty, R. F. et al. Temporal-callosal pathway diffusivity predicts phonological skills in children. Proc Natl Acad Sci U S A 104, 8556–8561, doi:10.1073/pnas.0608961104 (2007).

4 Frye, R. E. et al. Splenium microstructure is related to two dimensions of reading skill. Neuroreport 19, 1627–1631, doi:10.1097/WNR.0b013e328314b8ee (2008).

5 Carter, J. C. et al. A dual DTI approach to analyzing white matter in children with dyslexia. Psychiatry Res 172, 215–219, doi:10.1016/j.pscychresns.2008.09.005 (2009).

6 Odegard, T. N., Farris, E. A., Ring, J., McColl, R. & Black, J. Brain connectivity in nonreading impaired children and children diagnosed with developmental dyslexia. Neuropsychologia 47, 1972–1977, doi:10.1016/j.neuropsychologia.2009.03.009 (2009).

7 Rimrodt, S. L., Peterson, D. J., Denckla, M. B., Kaufmann, W. E. & Cutting, L. E. White matter microstructural differences linked to left perisylvian language network in children with dyslexia. Cortex 46, 739–749, doi:10.1016/j.cortex.2009.07.008 (2010).

8 Yeatman, J. D. et al. Anatomical properties of the arcuate fasciculus predict phonological and reading skills in children. J Cogn Neurosci 23, 3304–3317, doi: 10.1162/jocn_a_00061 (2011).

9 Zhang, M. et al. Language-general and -specific white matter microstructural bases for reading. Neuroimage 98, 435–441, doi:10.1016/j.neuroimage.2014.04.080 (2014).

10 Chiang, M. C. et al. Genetics of brain fiber architecture and intellectual performance. J Neurosci 29, 2212–2224, doi:10.1523/JNEUROSCI.4184-08.2009 (2009).

11 Budisavljevic, S. et al. Age-Related Differences and Heritability of the Perisylvian Language Networks. J Neurosci 35, 12625–12634, doi:10.1523/JNEUROSCI.1255-14.2015 (2015).

12 Powers, S. J., Wang, Y., Beach, S. D., Sideridis, G. D. & Gaab, N. Examining the relationship between home literacy environment and neural correlates of phonological processing in beginning readers with and without a familial risk for dyslexia: an fMRI study. Ann Dyslexia 66, 337–360, doi:10.1007/s11881-016-0134-2 (2016).

13 Vanderauwera, J., Wouters, J., Vandermosten, M. & Ghesquiere, P. Early dynamics of white matter deficits in children developing dyslexia. Dev Cogn Neurosci 27, 69–77, doi: 10.1016/j.dcn.2017.08.003 (2017).

14 Saygin, Z. M. et al. Tracking the roots of reading ability: white matter volume and integrity correlate with phonological awareness in prereading and early-reading kindergarten children. J Neurosci 33, 13251–13258, doi:10.1523/JNEUROSCI.4383-12.2013 (2013).

15 Langer, N. et al. White Matter Alterations in Infants at Risk for Developmental Dyslexia. Cereb Cortex 27, 1027–1036, doi:10.1093/cercor/bhv281 (2017).

16 Wang, Y. et al. Development of Tract-Specific White Matter Pathways During Early Reading Development in At-Risk Children and Typical Controls. Cereb Cortex 27, 2469–2485, doi:10.1093/cercor/bhw095 (2017).

17 Hoeft, F. et al. Neural systems predicting long-term outcome in dyslexia. Proc Natl Acad Sci U S A 108, 361–366, doi:10.1073/pnas.1008950108 (2011).

18 Myers, C. A. et al. White matter morphometric changes uniquely predict children’s reading acquisition. Psychol Sci 25, 1870–1883, doi:10.1177/0956797614544511 (2014).

19 Vandermosten, M. et al. A DTI tractography study in pre-readers at risk for dyslexia. Dev Cogn Neurosci 14, 8–15, doi:10.1016/j.dcn.2015.05.006 (2015).

20 Gabrieli, J. D., Ghosh, S. S. & Whitfield-Gabrieli, S. Prediction as a humanitarian and pragmatic contribution from human cognitive neuroscience. Neuron 85, 11–26, doi: 10.1016/j.neuron.2014.10.047 (2015).

21 Demerens, C. et al. Induction of myelination in the central nervous system by electrical activity. Proc Natl Acad Sci U S A 93, 9887–9892 (1996).

22 Stevens, B., Tanner, S. & Fields, R. D. Control of myelination by specific patterns of neural impulses. J Neurosci 18, 9303–9311 (1998).

23 Gibson, E. M. et al. Neuronal activity promotes oligodendrogenesis and adaptive myelination in the mammalian brain. Science 344, 1252304, doi:10.1126/science.1252304 (2014).

24 Hines, J. H., Ravanelli, A. M., Schwindt, R., Scott, E. K. & Appel, B. Neuronal activity biases axon selection for myelination in vivo. Nat Neurosci 18, 683–689, doi:10.1038/nn.3992 (2015).

25 Blumenfeld-Katzir, T., Pasternak, O., Dagan, M. & Assaf, Y. Diffusion MRI of structural brain plasticity induced by a learning and memory task. PLoS One 6, e20678, doi: 10.1371/journal.pone.0020678 (2011).

26 Sampaio-Baptista, C. et al. Motor skill learning induces changes in white matter microstructure and myelination. J Neurosci 33, 19499–19503, doi:10.1523/JNEUROSCI.3048-13.2013 (2013).

27 McKenzie, I. A. et al. Motor skill learning requires active central myelination. Science 346, 318–322, doi: 10.1126/science.1254960 (2014).

28 Bengtsson, S. L. et al. Extensive piano practicing has regionally specific effects on white matter development. Nat Neurosci 8, 1148–1150, doi:10.1038/nn1516 (2005).

29 Scholz, J., Klein, M. C., Behrens, T. E. & Johansen-Berg, H. Training induces changes in white-matter architecture. Nat Neurosci 12, 1370–1371 (2009).

30 Sagi, Y. et al. Learning in the fast lane: new insights into neuroplasticity. Neuron 73, 1195–1203, doi: 10.1016/j.neuron.2012.01.025 (2012).

31 Hofstetter, S., Tavor, I., Tzur Moryosef, S. & Assaf, Y. Short-term learning induces white matter plasticity in the fornix. J Neurosci 33, 12844–12850, doi:10.1523/JNEUROSCI.4520-12.2013 (2013).

32 Keller, T. A. & Just, M. A. Altering cortical connectivity: remediation-induced changes in the white matter of poor readers. Neuron 64, 624–631 (2009).

33 Gebauer, D. et al. Differences in integrity of white matter and changes with training in spelling impaired children: a diffusion tensor imaging study. Brain Struct Funct 217, 747–760, doi: 10.1007/s00429-011-0371-4 (2012).

34 Hofstetter, S., Friedmann, N. & Assaf, Y. Rapid language-related plasticity: microstructural changes in the cortex after a short session of new word learning. Brain Struct Funct 222, 1231–1241 doi: 10.1007/s00429-016-1273-2 (2017).

35 Vandermosten, M. et al. A tractography study in dyslexia: neuroanatomic correlates of orthographic, phonological and speech processing. Brain 135, 935–948, doi:10.1093/brain/awr363 (2012).

36 Gullick, M. M. & Booth, J. R. The direct segment of the arcuate fasciculus is predictive of longitudinal reading change. Dev Cogn Neurosci 13, 68–74, doi:10.1016/j.dcn.2015.05.002 (2015).

37 Wandell, B. A. & Yeatman, J. D. Biological development of reading circuits. Curr Opin Neurobiol 23, 261–268, doi:10.1016/j.conb.2012.12.005 (2013).

38 Yeatman, J. D., Dougherty, R. F., Ben-Shachar, M. & Wandell, B. A. Development of white matter and reading skills. Proc Natl Acad Sci U S A 109, E3045–3053, doi: 10.1073/pnas.1206792109 (2012).

39 Rauschecker, A. M. et al. Reading impairment in a patient with missing arcuate fasciculus. Neuropsychologia 47, 180–194, doi:10.1016/j.neuropsychologia.2008.08.011 (2009).

40 Catani, M., Jones, D. K. & ffytche, D. H. Perisylvian language networks of the human brain. Ann Neurol 57, 8–16, doi:10.1002/ana.20319 (2005).

41 Cohen, L. et al. Language-specific tuning of visual cortex? Functional properties of the Visual Word Form Area. Brain 125, 1054–1069 (2002).

42 Dehaene, S., Le Clec, H. G., Poline, J. B., Le Bihan, D. & Cohen, L. The visual word form area: a prelexical representation of visual words in the fusiform gyrus. Neuroreport 13, 321–325 (2002).

43 McCandliss, B. D., Cohen, L. & Dehaene, S. The visual word form area: expertise for reading in the fusiform gyrus. Trends Cogn Sci 7, 293–299 (2003).

44 Dehaene, S. & Cohen, L. The unique role of the visual word form area in reading. Trends Cogn Sci 15, 254–262, doi: 10.1016/j.tics.2011.04.003 (2011).

45 Bouhali, F. et al. Anatomical connections of the visual word form area. J Neurosci 34, 15402–15414, doi: 10.1523/JNEUROSCI.4918-13.2014 (2014).

46 Wandell, B. A., Rauschecker, A. M. & Yeatman, J. D. Learning to see words. Annu Rev Psychol 63, 31–53, doi:10.1146/annurev-psych-120710-100434 (2012).

47 Yeatman, J. D., Rauschecker, A. M. & Wandell, B. A. Anatomy of the visual word form area: adjacent cortical circuits and long-range white matter connections. Brain Lang 125, 146–155, doi: 10.1016/j.bandl.2012.04.010 (2013).

48 Rumsey, J. M. et al. Phonological and orthographic components of word recognition. A PET-rCBF study. Brain 120 (Pt 5), 739–759 (1997).

49 DeWitt, I. & Rauschecker, J. P. Wernicke’s area revisited: parallel streams and word processing. Brain Lang 127, 181–191 (2013).

50 Poldrack, R. A. et al. Functional specialization for semantic and phonological processing in the left inferior prefrontal cortex. Neuroimage 10, 15–35, doi: 10.1006/nimg.1999.0441 (1999).

51 Schrank, F. A., Mather, N., & McGrew, K. S. Woodcock-Johnson IV Tests of Achievement. (2014).

52 Torgesen, J., Wagner, R., Rashotte, C. Test of Word Reading Efficiency, Second Edition (TOWRE-2). (2012).

53 Deutsch, G. K. et al. Children’s reading performance is correlated with white matter structure measured by diffusion tensor imaging. Cortex 41, 354–363 (2005).

54 Lebel, C. & Beaulieu, C. Longitudinal development of human brain wiring continues from childhood into adulthood. J Neurosci 31, 10937–10947, doi:10.1523/JNEUROSCI.5302-10.2011 (2011).

55 Vandermosten, M., Boets, B., Wouters, J. & Ghesquiere, P. A qualitative and quantitative review of diffusion tensor imaging studies in reading and dyslexia. Neurosci Biobehav Rev 36, 1532–1552, doi: 10.1016/j.neubiorev.2012.04.002 (2012).

56 Lebel, C. et al. Diffusion tensor imaging correlates of reading ability in dysfluent and nonimpaired readers. Brain Lang 125, 215–222, doi: 10.1016/j.bandl.2012.10.009 (2013).

57 Horowitz-Kraus, T., Wang, Y., Plante, E. & Holland, S. K. Involvement of the right hemisphere in reading comprehension: a DTI study. Brain Res 1582, 34–44, doi: 10.1016/j.brainres.2014.05.034 (2014).

58 Horowitz-Kraus, T., Vannest, J. J., Gozdas, E. & Holland, S. K. Greater Utilization of Neural-Circuits Related to Executive Functions is Associated with Better Reading: A Longitudinal fMRI Study Using the Verb Generation Task. Front Hum Neurosci 8, 447, doi: 10.3389/fnhum.2014.00447 (2014).

59 Schwarz, G. Estimating the Dimension of a Model. The Annals of Statistics 6, 461–464 (1978).

60 Neath, A. A. C., J. E. The Bayesian information criterion: background, derivation, and applications. Wiley Interdiscip. Rev. Comput. Stat. 4, 199–203 (2012).

61 Yeatman, J. D., Dougherty, R. F., Myall, N. J., Wandell, B. A. & Feldman, H. M. Tract profiles of white matter properties: automating fiber-tract quantification. PLoS One 7, e49790, doi: 10.1371/journal.pone.0049790 (2012).

62 Mezer, A. et al. Quantifying the local tissue volume and composition in individual brains with magnetic resonance imaging. Nat Med 19, 1667–1672, doi: 10.1038/nm.3390 (2013).

63 Hedeker, D. & Gibbons, R. D. Longitudinal Data Analysis. (John Wiley & Sons, Inc., 2006).

64 Barquero, L. A., Davis, N. & Cutting, L. E. Neuroimaging of reading intervention: a systematic review and activation likelihood estimate meta-analysis. PLoS One 9, e83668, doi: 10.1371/journal.pone.0083668 (2014).

65 Boltzmann, M., Mohammadi, B., Samii, A., Munte, T. F. & Russeler, J. Structural changes in functionally illiterate adults after intensive training. Neuroscience 344, 229–242, doi: 10.1016/j.neuroscience.2016.12.049 (2017).

66 Kanai, R. & Rees, G. The structural basis of inter-individual differences in human behaviour and cognition. Nat Rev Neurosci 12, 231–242, doi:10.1038/nrn3000 (2011).

67 Tsang, J. M., Dougherty, R. F., Deutsch, G. K., Wandell, B. A. & Ben-Shachar, M. Frontoparietal white matter diffusion properties predict mental arithmetic skills in children. Proc Natl Acad Sci U S A 106, 22546–22551, doi:10.1073/pnas.0906094106 (2009).

68 Van Beek, L., Ghesquiere, P., Lagae, L. & De Smedt, B. Left fronto-parietal white matter correlates with individual differences in children’s ability to solve additions and multiplications: a tractography study. Neuroimage 90, 117–127, doi: 10.1016/j.neuroimage.2013.12.030 (2014).

69 van Eimeren, L., Niogi, S. N., McCandliss, B. D., Holloway, I. D. & Ansari, D. White matter microstructures underlying mathematical abilities in children. Neuroreport 19, 1117–1121, doi: 10.1097/WNR.0b013e328307f5c1 (2008).

70 Matejko, A. A., Price, G. R., Mazzocco, M. M. & Ansari, D. Individual differences in left parietal white matter predict math scores on the Preliminary Scholastic Aptitude Test. Neuroimage 66, 604–610, doi:10.1016/j.neuroimage.2012.10.045 (2013).

71 Navas-Sanchez, F. J. et al. White matter microstructure correlates of mathematical giftedness and intelligence quotient. Hum Brain Mapp 35, 2619–2631, doi:10.1002/hbm.22355 (2014).

72 Jolles, D. et al. Plasticity of left perisylvian white-matter tracts is associated with individual differences in math learning. Brain Struct Funct 221, 1337–1351, doi:10.1007/s00429-014-0975-6 (2016).

73 Sykova, E. & Nicholson, C. Diffusion in brain extracellular space. Physiol Rev 88, 1277–1340, doi: 10.1152/physrev.00027.2007 (2008).

74 Walhovd, K. B., Johansen-Berg, H. & Karadottir, R. T. Unraveling the secrets of white matter-bridging the gap between cellular, animal and human imaging studies. Neuroscience 276, 2–13, doi: 10.1016/j.neuroscience.2014.06.058 (2014).

75 Lutti, A., Dick, F., Sereno, M. I. & Weiskopf, N. Using high-resolution quantitative mapping of R1 as an index of cortical myelination. Neuroimage 93 Pt 2, 176–188, doi: 10.1016/j.neuroimage.2013.06.005 (2014).

76 Yeatman, J. D., Wandell, B. A. & Mezer, A. A. Lifespan maturation and degeneration of human brain white matter. Nat Commun 5, 4932, doi:10.1038/ncomms5932 (2014).

77 Weiskopf, N., Mohammadi, S., Lutti, A. & Callaghan, M. F. Advances in MRI-based computational neuroanatomy: from morphometry to in-vivo histology. Curr Opin Neurol 28, 313–322, doi: 10.1097/WCO.0000000000000222 (2015).

78 Travis, K. E., Ben-Shachar, M., Myall, N. J. & Feldman, H. M. Variations in the neurobiology of reading in children and adolescents born full term and preterm. Neuroimage Clin 11, 555–565, doi: 10.1016/j.nicl.2016.04.003 (2016).

79 Duara, R. et al. Neuroanatomic differences between dyslexic and normal readers on magnetic resonance imaging scans. Arch Neurol 48, 410–416 (1991).

80 Rumsey, J. M. et al. Corpus callosum morphology, as measured with MRI, in dyslexic men. Biol Psychiatry 39, 769–775 (1996).

81 von Plessen, K. et al. Less developed corpus callosum in dyslexic subjects--a structural MRI study. Neuropsychologia 40, 1035–1044 (2002).

82 Bell, N. Seeing stars. (Gander, 2007).

83 Caruyer, E., Aganj, I., Lenglet, C., Sapiro, G. & Deriche, R. Motion Detection in Diffusion MRI via Online ODF Estimation. Int J Biomed Imaging 2013, 849363, doi: 10.1155/2013/849363 (2013).

84 Andersson, J. L., Skare, S. & Ashburner, J. How to correct susceptibility distortions in spin-echo echo-planar images: application to diffusion tensor imaging. Neuroimage 20, 870–888, doi: 10.1016/S1053-8119(03)00336-7 (2003).

85 Andersson, J. L. & Sotiropoulos, S. N. An integrated approach to correction for off-resonance effects and subject movement in diffusion MR imaging. Neuroimage 125, 1063–1078, doi: 10.1016/j.neuroimage.2015.10.019 (2016).

86 Jensen, J. H., Helpern, J. A., Ramani, A., Lu, H. & Kaczynski, K. Diffusional kurtosis imaging: the quantification of non-gaussian water diffusion by means of magnetic resonance imaging. Magn Reson Med 53, 1432–1440, doi:10.1002/mrm.20508 (2005).

87 Garyfallidis, E. et al. Dipy, a library for the analysis of diffusion MRI data. Front Neuroinform 8, 8, doi:103389/fninf2.01400008 (2014).

88 Yendiki, A., Koldewyn, K., Kakunoori, S., Kanwisher, N. & Fischl, B. Spurious group differences due to head motion in a diffusion MRI study. Neuroimage 88, 79–90, doi: 10.1016/j.neuroimage.2013.11.027 (2014).

89 Tournier, J. D., Calamante, F., Gadian, D. G. & Connelly, A. Direct estimation of the fiber orientation density function from diffusion-weighted MRI data using spherical deconvolution. Neuroimage 23, 1176–1185, doi:10.1016/j.neuroimage.2004.07.037 (2004).

